# β-strand complementation within tip initiation complexes licenses assembly of diverse type IV filaments

**DOI:** 10.64898/2026.02.14.705861

**Authors:** Taylor J. Ellison, Zoey R. Litt, Justin N. Applegate, Emma Miller, Luther Davis, Alfredo Ruiz-Rivera, Angela Lisovsky, Rittika Saha, Naveen Jasti, Stephanie McLaughlin, Beth Traxler, Courtney K. Ellison, Alexey J. Merz

## Abstract

PilC/PilY1 proteins are tip-located adhesins of type IV pili (T4P) that are critical for T4P function in diverse behaviors including twitching motility, DNA uptake, and host cell adhesion. PilC and PilY1 adhesins are proposed to interact with initiation complexes composed of minor pilins (PilIJK family proteins) to aid in initiation of T4P polymerization, but it has been unclear how PilC/PilY1 proteins promote fiber assembly. We combined structural modeling, genetic, and biochemical experiments using *Neisseria gonorrhoeae* and *Acinetobacter baylyi* to delineate how PilC/PilY1 control T4P assembly: a short peptide at the C-terminus of PilC/PilY1 initiates T4P assembly via β-strand complementation with PilK-family minor pilins. This β-strand is necessary and partially sufficient to trigger fiber assembly. In a working model, the PilK-PilC/PilY1 complex is recognized by a preformed PilI-PilJ heterodimer to form a quaternary “licensing complex” that then templates and initiates fiber assembly. In type II secretion systems (T2SS) lacking PilC/PilY1, PilK homologs directly incorporate the terminal β-strand provided by PilC/PilY1 in T4P. Moreover, phylogenetically distinct Tad T4P lack a canonical PilK homolog and instead contain a structurally similar minor pilin-like protein called TadG/CpaL that is important for fiber assembly. We show that CpaL of *Caulobacter crescentus* Tad T4P acts similarly to the T2SS PilK homolog to provide the C-terminal β-strand required for assembly. Our results explain how PilC/PilY1 can be retained on the fiber tip under enormous tensile loads generated during mechanical shear and T4P retraction and demonstrate how diverse T4P systems employ β-strand complementation to license fiber assembly.

**SIGNIFICANCE:** Prokaryotic type IV filaments are ancient, diverse, and broadly distributed nanomachines that assemble and retract to execute diverse microbial functions. They include type IV pili and type II secretion systems, mediating toxin secretion, motility, surface adhesion, biofilm formation, DNA uptake, and other functions. Here, we show that two widely conserved subunits of the tip, PilI and PilJ, form a module that recognizes the folding of a β-sheet in a third subunit, PilK. The final β-strand in this sheet can be supplied *in trans* by the last ∼10 aminoacyl residues of large PilC/PilY1 adhesins, or *in cis* by PilK itself. In a working model, this recognition results in formation of a PilIJK trimer, which then licenses fiber polymerization through a templating mechanism.

## INTRODUCTION

The type IV filaments (T4F) of Bacteria and Archaea are related, dynamic machines that reversibly assemble filamentous polymers to execute diverse functions including surface attachment and motility, DNA uptake, virulence, and protein secretion (1, 2). The T4F superfamily comprises diverse microbial appendages including type IV pili (T4P), mannose-sensitive hemagglutinin (MSHA) pili, type II secretion systems (T2SS), and the archaellum used for archaeal swimming motility (3, 4). Bacterial T4P can further be subdivided: T4aP, T4bP, and Tad T4cP are distinguished by differences in pilin subunit leader sequence length and homologies among assembly components (4). The competence (Com) pili of Gram-positive species form extended filaments and are now classified as T4dP (5–9). Most T4F share the ability to dynamically extend and retract. T4F dynamics are driven by cyclical polymerization and disassembly of pilin subunits from a cytoplasmic membrane pool (10–16). While extension and retraction dynamics are crucial for most T4F functions, archaella instead rotate to power swimming, similar to bacterial flagella.

Pilin subunits are added and removed from the fiber base. Initiation relies on minor pilin initiation complexes at the filament apical tip. However, the assembly sequences and quaternary structures of these complexes are opaque (17–20). T4P tip complexes commonly comprise three or more distinct minor pilin subunits, each structurally similar to the major pilin that comprises the bulk of the fiber. Pilins fold into a “lollipop” structure with a conserved N-terminal hydrophobic domain and a C-terminal globular domain that is structurally more variable. The N-terminal α-helical domain anchors unassembled pilins in the cytoplasmic membrane and forms the structural core of the polymerized fiber. The globular C-terminal domain is more exposed and contributes to the functional diversity of the T4F superfamily. Polymer initiation can occur 200 times per minute per cell (21). Minor pilin initiation complexes are thought to function analogously to eukaryotic Arp2/3 and γ-tubulin ring complexes that initiate actin filament and microtubule polymerization (22, 23). In each case, an oligomer of subunits homologous to the major protomer is hypothesized to nucleate polymerization through a templating mechanism. Templating increases the rate of polymer initiation by overcoming rate-limiting thermodynamic barriers that impede spontaneous nucleation. Arp2/3 and γ-tubulin complexes further act as signal integrators that license filament polymerization in response to specific cues. T4P tip complexes may have analogous functions.

In addition to minor pilins, T4aP tips often incorporate large adhesins of the PilC/PilY1 family (19, 24–27). In mutants lacking individual minor pilins or PilC/PilY1, many species including *Neisseria gonorrhoeae*, *Pseudomonas aeruginosa*, *Acinetobacter baylyi*, and *Myxococcus xanthus* are nearly devoid of pili at steady state. Removal of PilT, the retraction motor ATPase that powers depolymerization of the pilus fiber, traps pili in the assembled state and suppresses these steady-state defects in piliation (10, 12, 17, 26, 28). An obvious inference is that minor pilins and PilC/PilY1 are required for efficient polymer initiation, but dispensable for processive filament polymerization or depolymerization.

PilC/PilY1 translocate across the cytoplasmic membrane through the SecYE/signal peptidase I pathway. They have structurally diverse N-terminal adhesin domains but share a conserved C-terminal β-propellor domain (29). They usually contain several disulfide bonds. N-terminal adhesin domains can be over 3,000 residues long, and are hypothesized to encompass diverse adhesin activities. Some evidence indicates that the C-terminal β-propellor is sufficient to support pilus assembly (30), and recent structural results prompted speculation that PilC/PilY1 has a role in gating of the outer membrane secretin bushing, through which the assembled fiber passes (Guo et al. Nat Comms 2024). However, the structure of the tip complex and the molecular mechanism(s) through which PilC/PilY1 controls fiber initiation have remained unclear. We now present molecular modeling, genetic, and biochemical evidence that a ∼6-10 aa assembly signal at the extreme C-terminus of PilC/PilY1 (Cβ) licenses formation of the core minor pilin trimer PilIJK through a β-strand complementation with PilK-family minor pilins. The PilK-PilC/PilY1 heterodimer is then recognized by a pre-formed PilI-PilJ heterodimer. Licensing by Cβ is necessary, and PilK-Cβ fusions are at least partially sufficient, to initiate fiber assembly and elongation. Structural modeling suggests that diverse T4F use this β-strand complementation for tip complex formation. In line with this prediction, we show the minor pilin-like protein CpaL found in Tad T4cP behaves similarly to GspK in T2SSs, providing the β-strand required for complex assembly. To our knowledge, this is the first example of a specific polymerization signal for any T4F system.

## RESULTS

Initiation tip complex protein nomenclature varies across T4F systems. For example, the *N. gonorrhoeae* minor pilins PilH, PilI, PilJ, and PilK correspond to FimT/U, PilV, PilW, and PilX in *P. aeruginosa*. We use existing annotations (except for the PilW/GspJ protein ComB in *A. baylyi*, here referred to as PilW). The names of homologous subunits in different T4F systems are shown in **Table 1 and Fig. S1.**

**Table 1.**
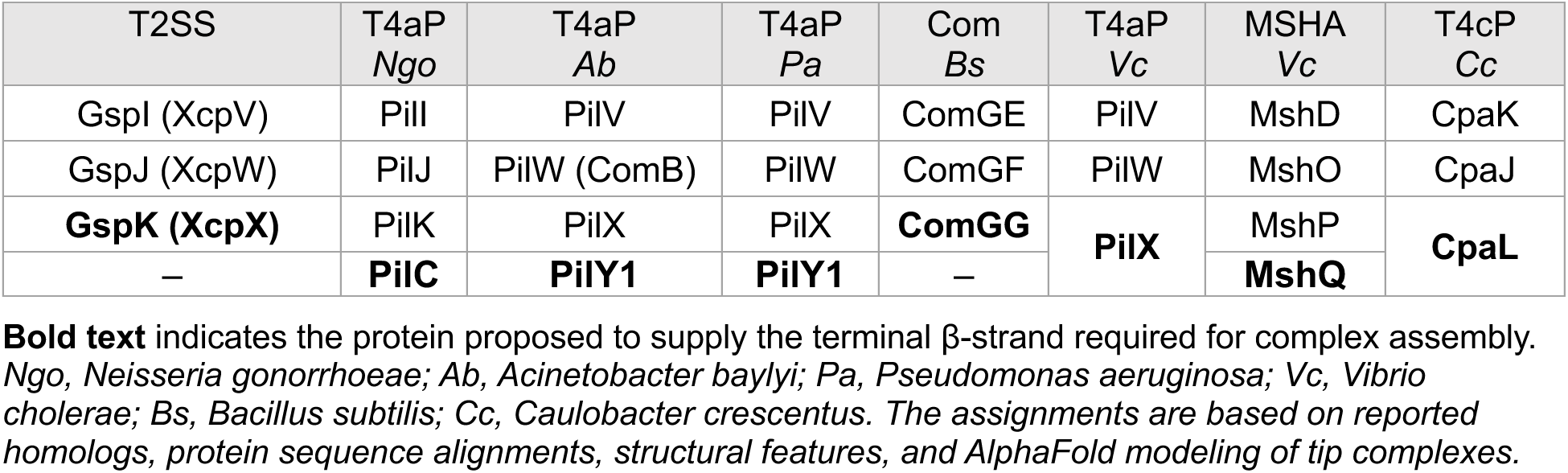
Core initiation tip complex homologs across different species and T4F systems.

### A PilC/PilY1 C-terminal peptide is predicted to strand-complement the β-sheet in GspK homologs

AlphaFold3 modeling of PilC/PilY1 complexes found in T4aP with core minor pilins yielded consistent predictions for the arrangement of PilI, PilJ, PilK, and PilC/PilY1 across diverse species including: *Neisseria gonorrhoeae; Acinetobacter baylyi*; *Pseudomonas aeruginosa*; and *Myxococcus xanthus*. A model of the *N. gonorrhoeae* tip complex is shown in **Fig. 1**. Additional T4aP models are shown in **Fig. S2**. **Figs. S3-S6** show confidence metrics for the structural predictions. In every case, PilI, PilJ, and PilK are arranged in clockwise order when viewed from above the complex’s apical tip (**Fig. 1A**). This predicted structural conservation parallels observations that the polycistronic organization of *pilI*, *pilJ*, and *pilK* genes is almost invariably syntenic across T4P and T2SS (17) (**Fig. S1**).

**Figure 1.**
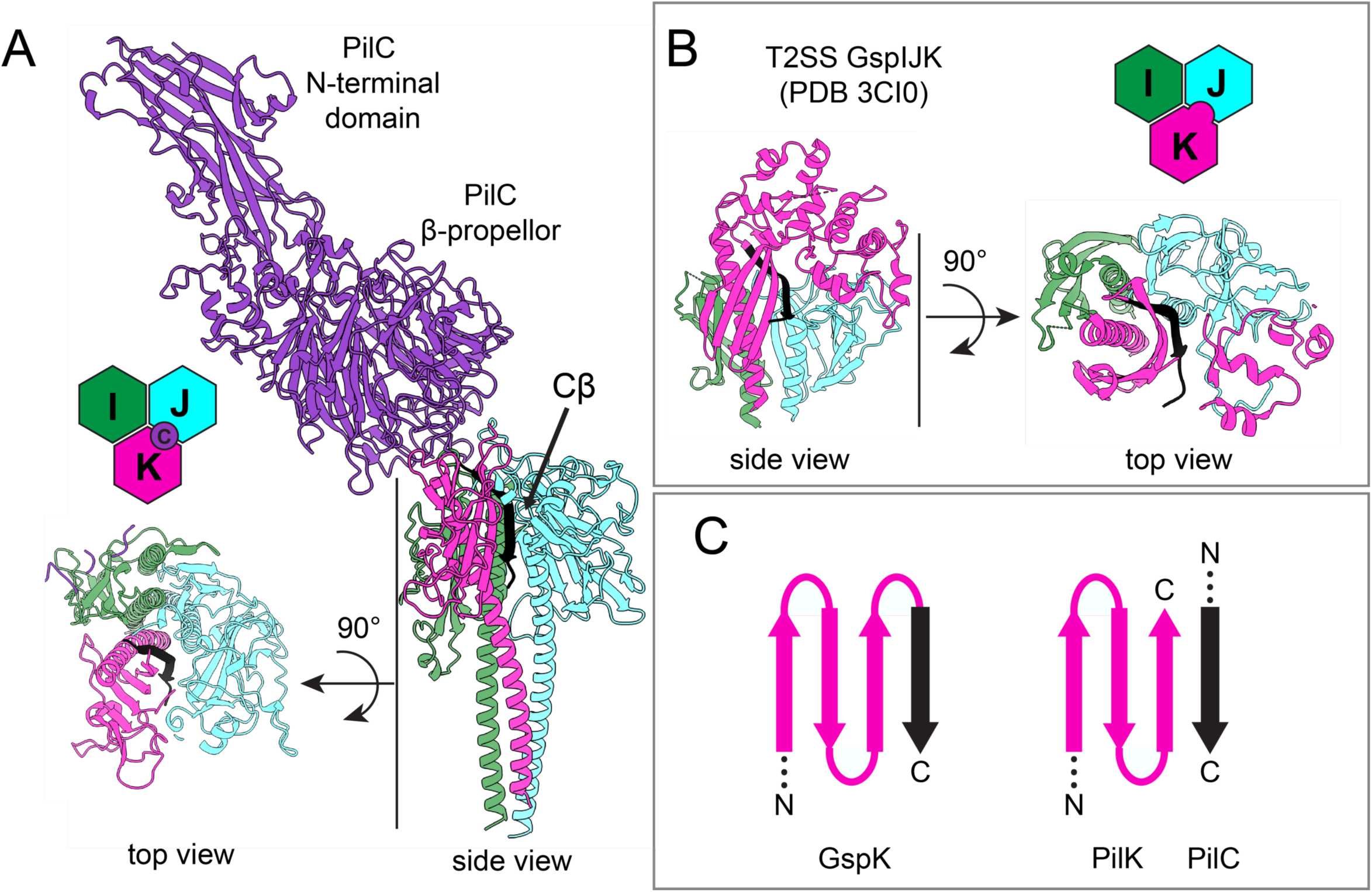
A conserved β-strand is predicted to strand-complement the β-sheet in GspK homologs. (A) AlphaFold-predicted model of the *N. gonorrhoeae* PilIJK minor pilins in complex with PilC. In *N. gonorrhoeae* the complementing β-strand is provided by PilC. (B) Structure of the T2SS minor pilin complex (PDB 3CI0) from enterotoxigenic *E. coli*. In T2SSs, the complementing β-strand is provided by GspK. (C) Schematic of the β-strand complementation model. In all figure panels, the conserved C-terminal β-strand is colored black.

The conserved PilC/PilY1 β-propellor is predicted to sit atop the minor pilin PilIJK heterotrimer, with variable orientations in different models. However, in every case the extreme C-terminus of PilC/PilY1 (∼10 aminoacyl residues) penetrates down the centerline of the PilI-J-K heterotrimer at the PilJ-PilK assembly interface (**Fig. 1A, Fig. S2**). In each case the PilC/PilY1 C-terminal peptide is predicted to strand-complement the small β-sheet in PilK, and lies in a complementary groove in PilJ (**Fig. 1A**). Solved crystal structures of the T2SS tip complexes of *Escherichia coli* (GspIJK) and *Pseudomonas aeruginosa* (XcpVWX) (31, 32) topologically mirror the predicted T4P tip complex. Remarkably, in the T2SS structures the conserved C-terminal β-strand is provided by PilK homologs GspK and XcpX, rather than by a tip adhesin (**Fig. 1B**).

This structural arrangement suggests a working model: stepwise association of the C-terminal PilC/PilY1 β-strand with the beta sheet in the globular domain of PilK is followed by recognition of this heterodimer by PilJ and PilI. Deep burial of the PilC/PilY1 C-terminal β-strand appears to rule out models in which PilC/PilY1 associates with a pre-formed PilI-J-K heterotrimer. The large buried area of the PilC β-strand in the predicted *N. gonorrhoeae* PilI-J-K complex (∼1700 Å^2^ for the example in **Fig. 1A**) would provide mechanical stability required to withstand enormous tensile forces (≥100 pN) developed during pilus retraction or when cells encounter hydrodynamic shear forces (12, 33).

### PilK and PilC, and PilI and PilJ, form heterodimers

PilK and its homologs are predicted to be the most apical minor pilins in the tip complex, as they invariably lack a conserved +5 Glu residue critical for interactions with the N-terminal regions of previously incorporated (more apical) pilin subunits. Coupled with the prediction that the PilC/PilY1 β-strand mimics the C-terminal β strand of GspK in T2SS (which lack PilC/PilY1), we hypothesize that PilC/PilY1 and PilK form a heterodimer that assembles prior to PilI-PilJ incorporation. To test whether PilK and PilC/PilY1 can form a heterodimeric structure, we co-expressed periplasmic, soluble *N. gonorrhoeae* PilK-His_6_ (34) and the conserved C-terminal half of PilC, bearing a cleavable N-terminal signal peptide and a FLAG tag (FLAG-ΔN-PilC) in *E. coli* cells. SDS-PAGE followed by silver stain of periplasmic fractions revealed co-elution of PilK and PilC, which were verified by immunoblot (**Fig. 2A**). Thus, as predicted by our modeling, PilK directly binds to the C-terminal half of PilC.

**Figure 2.**
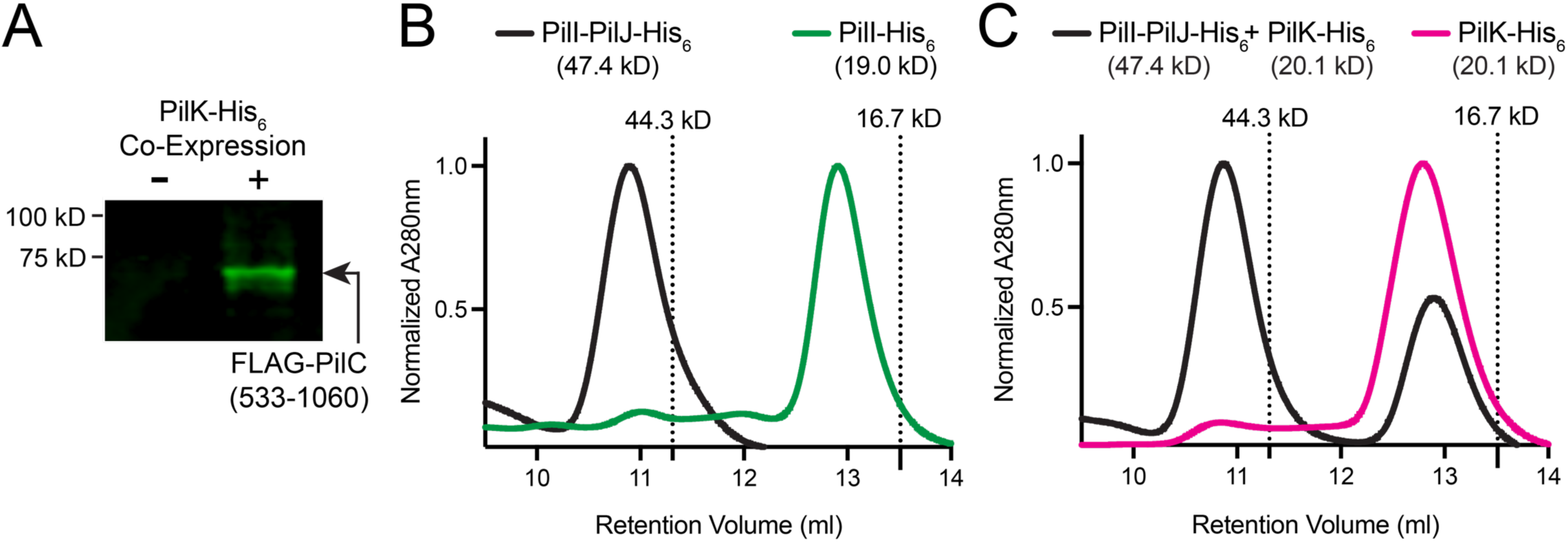
PilK and PilC, and PilI and PilJ, form heterodimers. (A) Co-Purification of periplasmically expressed FLAG-PilC(533-1060) when PilK-His_6_ is captured by immobilized metal chromatography. Eluates were immunoblotted with anti-FLAG antibodies to detect FLAG-tagged PilC. FLAG-PilC(533-1060) was expressed in the absence (lane 1) or presence (lane 2) of PilK-His_6_. (B) Size exclusion chromatography (Superdex 75 Increase) of soluble PilI-PilJ-His_6_ heterodimer (black) or PilI-His_6_ alone (green). (C) Size exclusion chromatography (Superdex 75 Increase) of purified soluble PilI-PilJ-His_6_ and PilK-His_6_ proteins mixed together (black) and PilK-His_6_ alone (pink). In figures (B) and (C), A280nm values are normalized to the maximum peak of each trace. Expected masses of the protein complexes are indicated in parentheses. Dashed lines indicate elution peaks for standards: ovalbumin (44.3 kDa) myoglobin (16.7 kDa). Each trace is representative of 3 independent replicates.

The T2SS GspI and GspJ minor pilins are adjacent in crystal structures (31, 32, 35). In our experiments, *N. gonorrhoeae* T4aP minor pilins PilI and PilJ formed a heterodimer stable through affinity co-purification (34) and size exclusion chromatography (**Fig. 2B**). Notably, addition of PilK to the heterodimeric sample did not result in a shift in the PilI-PilJ elution curve (**Fig. 2C**). In *N. gonorrhoeae*, deletion of either PilI or PilJ results in loss of both proteins, without affecting the stability of other minor pilins or PilC (17). Moreover, PilJ alone could not be purified, while co-expression with PilI allowed PilJ purification (34). Together these results indicate that PilI and PilJ interact to form a stable, obligate heterodimer. We therefore propose that a pre-assembled PilI-PilJ heterodimer in turn recognizes formation of the PilC/PilY1 complex. PilC/PilY1 proteins are large. *A. baylyi* PilY1 is ∼1500 residues long and has ten predicted disulfide bonds. Metagenomic sequences identify PilC/PilY1 family members of up to 3500 residues. We surmise that cells must ensure that PilC/PilY1 is synthesized, translocated into the periplasm, and folded before being mounted at the tip of a rapidly polymerizing T4P. Licensing of initiation complex assembly by the extreme C-terminus of PilC/PilY1 would satisfy this requirement.

### The C-terminal peptide is essential and partially sufficient for PilC function

Most *N. gonorrhoeae* strains encode two redundant and phase-variable PilC/PilY1-encoding genes. They appear to be functionally equivalent (24, 36). In strain FA1090 *pilC1* is phase-varied on, while *pilC2* is phase-varied off. To simplify our experiments, we employed a Δ*pilC2* mutant background, with *pilC1* phase-locked on (see strain table for details). Mutant cells lacking the 12-residue C-terminal peptide of PilC1 (*pilC1*Δ*Cβ*) phenocopied the Δ*pilC1* null mutant. Consistent with our hypothesis that the PilC/PilY1 β-strand is required for minor pilin complex licensing, T4P-labeling using cysteine knock-in maleimide-staining (21, 37, 38) revealed loss of fiber assembly and dynamics in Δ*pilC1* and *pilC1*Δ*Cβ* mutants (**Fig. 3A, B; Video SV1**). Similarly, colony morphology resembled that of mutants lacking T4P (**Fig. S7**). Natural transformation, which depends on T4P-mediated DNA uptake, was also abolished in the *pilC1*Δ*Cβ* mutant (**Fig. 3C**). The *pilC1*Δ*Cβ* allele exhibits partially dominant loss of function, so restoration of function *pilC1*Δ*Cβ* mutant cells through complementation *in trans* was not possible. Thus, we turned to amber suppression. Insertion of an amber stop codon *pilC1-Lys1051Amb* to truncate *pilC1* phenocopied the loss of piliation and DNA transformation caused by *pilC1*Δ*β* Expression of *tRNA^Lys^_Amb_* rescued the DNA transformation defect *Lys1051Amb* by about 20-fold (**Fig. 3D**). To our knowledge this is the first use of tRNA suppression in *Neisseriaceae*. Future work should result in systems capable of more efficient suppression, as well as incorporation of noncanonical amino acids.

**Figure 3.**
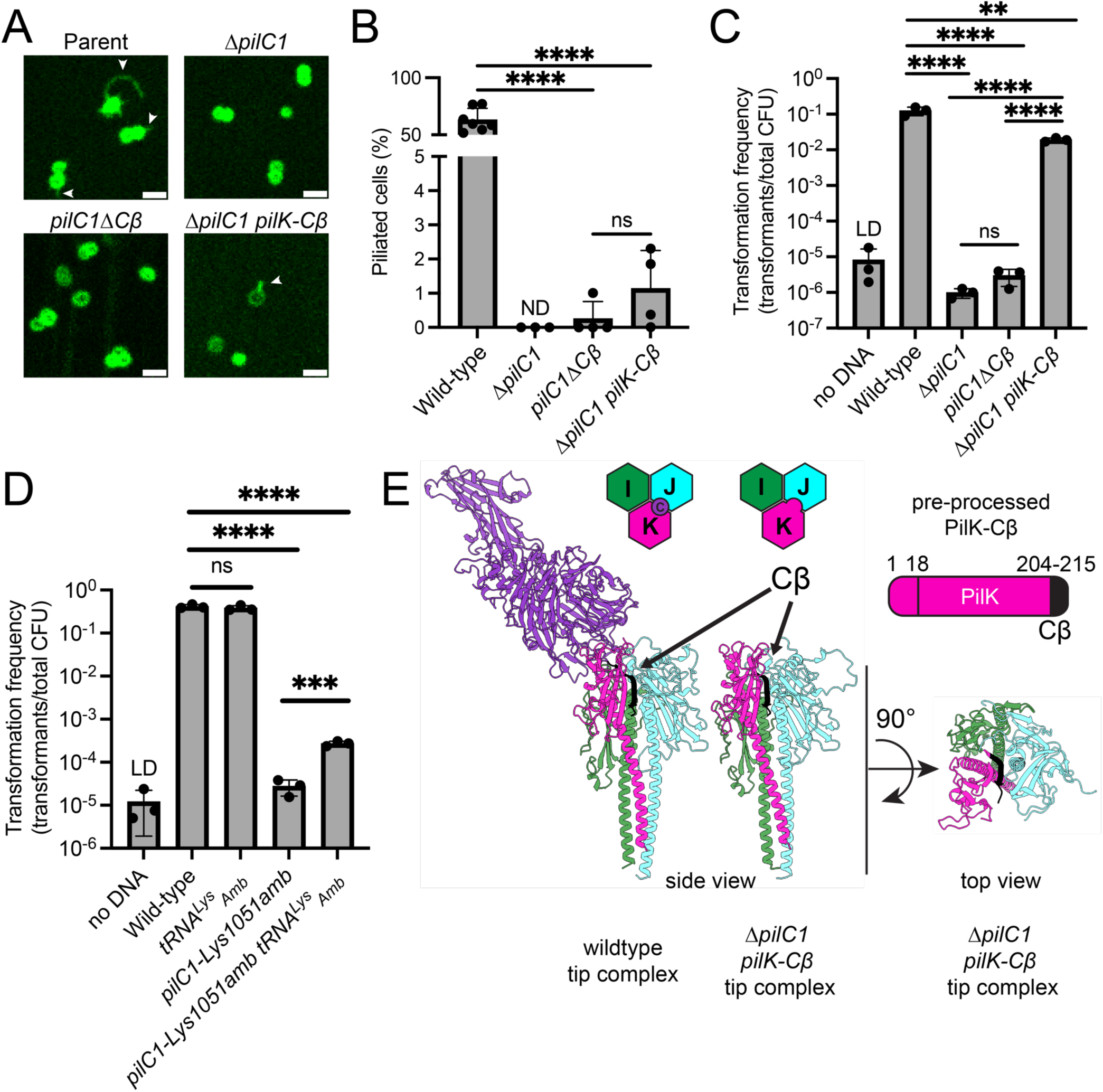
The extreme C-terminus of PilC is essential and partially sufficient for its function in *N. gonorrhoeae*. (A) Representative microscopy images of indicated strains. White arrows indicate examples of visible T4P filaments. Scale bars, 2 µm. (B) Quantification of piliated cells in populations of strains shown in (A). Bar graph shows the mean +/- SD. Each dot represents an independent, biological replicate. ND, not detected. (C and D) Natural transformation assay data from indicated strains. Bar graphs shows the mean +/- SD. Each dot represents an independent, biological replicate. (E) AlphaFold models of T4P tip complexes from indicated strains. Top right shows a schematic of the pre-processed form of the PilK-Cβ fusion. Statistical analysis for natural transformation assays was performed on log-transformed transformation frequency data. Statistics were determined by one-way ANOVA and corrected by Sidak’s multiple comparisons test. ****P<0.0001; **P<0.005; ns, not significant; LD, limit of detection.

In crystal structures of the T2SS PilK ortholog GspK, an additional C-terminal β-strand is present, at the location where the last ten residues of PilC/PilY1 are predicted to engage in β-strand complementation with PilK (**Fig. 1**). We therefore speculated that fusion of the last ∼10-20 residues of PilC to the C-terminus of the minor pilin PilK might compensate for loss of PilC/PilY1. AlphaFold modeling confidently predicted a fusion of the terminal residues from PilC to the C-terminus of PilK could form a stable complex, with the fused C-terminal β-strand positioned between PilK and PilJ in the same location as the wild-type PilC C-terminal β-strand (**Fig. 3E**). We constructed a Δ*pilC1* strain where the terminal 12 residues from PilC1 were appended to the extreme C-terminus of PilK (annotated here as *pilK-Cβ*) (**Fig. 3E**). While the Δ*pilC1* mutant produced no visible T4P, and T4P were only observed in a single replicate of the *pilC1*Δ*Cβ* strain experiments, T4P were apparent in 3 of 4 biological replicates for the *pilK-Cβ* strain suggesting an increase in piliation versus the *pilC1* mutants (**Fig. 3A, B**). Although this difference was not statistically significant, long pili are relatively rare and the limit of detection by fluorescence microscopy is <250 nm, limiting detection of shorter fibers (21). To test whether the *pilK-Cβ* strain might produce an increase in functional but short T4P, we performed natural transformation assays. The *pilK-Cβ* allele increased transformation frequency by more than four orders of magnitude versus the Δ*pilC1* null allele, reaching almost wild-type efficiency (**Fig. 3C**). Thus, the last 12 residues of PilC are partially sufficient for pilus initiation and function when full-length PilC is completely absent.

### The PilC/PilY1 β-strand’s function is conserved

To determine whether the C-terminal β-strand of PilC1/PilY1 functions similarly in other T4aP systems, we used *Acinetobacter baylyi* as a model organism to represent the well-studied T4P found in the *Pseudomonadales* order, which includes *P. aeruginosa* and closely related *Acinetobacter* pathogens. Similar to *N. gonorrhoeae* T4aP, AlphaFold modeling predicted that the extreme C-terminal residues of PilY1 form a β-strand that completes the β-sheet of PilX, the *A. baylyi* PilK homolog (**Fig. 4A, S2, S4**). As in *N. gonorrhoeae*, C-terminal β-strand deletion from PilY1 in *A. baylyi* phenocopied the Δ*pilY1* deletion and caused loss of pilus assembly (**Fig. 4B, C**). In *A. baylyi*, we observed a significant increase in piliation in a Δ*pilY1 pilX*-*Cβ* strain (**Fig 4C**). Natural transformation experiments also indicated restoration of pilus function, with the Δ*pilY1 pilX*-*Cβ* strain showing a ∼10^4^-fold increase over the Δ*pilY1* strain, exactly mirroring our experiments with *N. gonorrhoeae*. (**Fig. 4D**). Together, our data demonstrate that the final ∼10 residues of PilC/PilY1 are necessary and partially sufficient for tip complex function in T4aP, in both *N. gonorrhoeae* and *A. Baylyii*.

**Figure 4.**
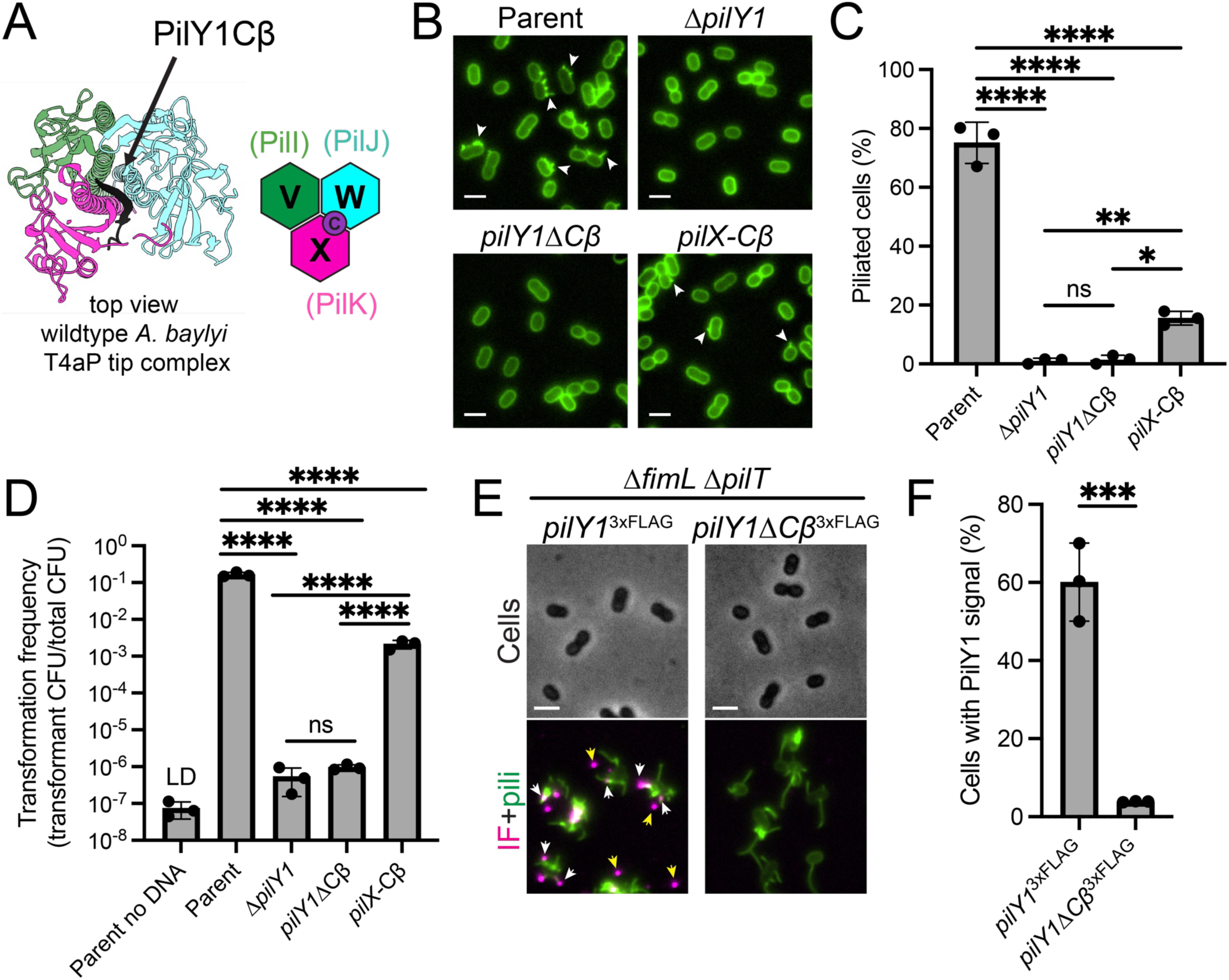
The extreme C-terminus of PilY1 is essential and partially sufficient for its function in other T4aP systems. (A) Top view of the AlphaFold model of the wildtype T4P tip complex from *A. baylyi*. (B) Representative microscopy images of indicated strains with background fluorescence subtracted. White arrows indicate examples of visible T4P filaments. (C) Quantification of piliated cells in populations of strains shown in (B). Bar graph shows the mean +/- SD. Each dot represents an independent, biological replicate. (D) Natural transformation assay data from indicated strains. Bar graph shows the mean +/- SD. Each dot represents an independent, biological replicate. (E) Representative microscopy images of indicated strains with background fluorescence subtracted. IF, immunofluorescence signal. White arrows represent examples of PilY1 associated with individual cells. Yellow arrows indicate sheared PilY1 that is not associated with cells. (F) Percentage of cells with associated PilY1 immunofluorescence signal of strains shown in (E). Bar graph shows the mean +/- SD. Each dot represents an independent, biological replicate. Statistical analysis for natural transformation assays was performed on log-transformed transformation frequency data. Statistics were determined by one-way ANOVA and corrected by Sidak’s multiple comparisons test. ****P<0.0001; ***P<0.001, **P<0.01; *P<0.05; ns, not significant; LD, limit of detection. Scale bars, 2 µm.

### The terminal β-strand stabilizes PilC/PilY1 at the filament tip

Because the C-terminal β-strand is predicted to anchor PilC/PilY1 to minor pilins in the tip complex, we reasoned that deletion of this peptide should prevent PilY1 association with the pilus tip. Using recently published immunofluorescence microscopy methods for labeling tip-associated minor pilins (39, 40), we tagged PilY1 and PilY1ΔCβ at the native locus with a 3xFLAG epitope and assessed whether cells exhibited associated PilY1 immunofluorescence signal (**Fig 4E, F**). Although the 3xFLAG tag caused a ∼1-log_10_ loss of function in natural transformation experiments, PilY1ΔCβ was present at similar levels as PilY1, showing that the C-terminal β-strand is not needed for PilY1 expression or stability *per se* (**Fig. S8A, B**). Because *pilY1*Δ*Cβ* mutant cells do not produce detectable T4P in cells where the PilT retraction ATPase is intact, we employed Δ*pilT* mutants to increase the amount of visible T4P for immunofluorescence studies. Additionally, *A. baylyi* produces its T4P close together in a line along the long axis of the cell making it difficult to resolve individual T4P, especially in hyperpiliated Δ*pilT* strains. Long-axis localization depends on the chemosensory Pil-Chp pathway (41, 42), so we used a published Pil-Chp mutation (Δ*fimL*) to enhance detection of individual, extended T4P (**Fig. 4E, F**). Wild-type PilY1 associated with ∼60% of cells versus ∼4% of cells expressing PilY1ΔCβ, indicating that the C-terminal β-strand is important for PilY1 association with the T4P fiber (**Fig. 4E, F**). While we expect that all T4P tip complexes in the wild type incorporate PilY1, not all cells or T4P exhibited detectable PilY1 signal, as in minor pilin immunofluorescence studies (39). This is likely due to shearing of T4P during the many wash steps in immunofluorescence sample preparation. Indeed, we frequently observed PilY1 signal not associated with cells, including sheared filaments with tip-associated PilY1 (**Fig. 4E, S8C**).

### Diverse T4Fs employ C-terminal β-strand complementation to stabilize tip complexes

T4P are broadly distributed structures that belong to the diverse T4F family composed of structurally related nanomachines. Detailed phylogenetic analysis of T4P and their homologs by Denise et al. 2019 shows T4F are so widely distributed that they may have been present in the last universal common ancestor (4, 13) (**Fig. S9**). Evolution of the T4F family has led to diversification of these structures into multiple distinct subclades, some with dedicated functions as exemplified by Com T4dP found in competent Gram-positive bacteria that are used specifically for DNA uptake (**Fig. S9**). Crystal structures and models of minor pilin trimers from T2SS and structural predictions of the tip complexes from other T4F provide support that β-strand complementation may promote tip complex licensing and stability in diverse systems (**Fig. 1B, Fig. S10**). In systems where minor pilin homologs are present and readily identified (see Discussion for details), a C-terminal β-strand is present at the same location and orientation as the predicted PilC/PilY1 C-terminal β-strand in T4aP (see **Fig. S11-15** for AlphaFold confidence metrics). The MSHA-encoding locus found in *V. cholerae* is not predicted to contain a PilY1 homolog, but the gene encoding the large (∼1200 residue) protein MshQ is located immediately downstream of the *pilIJK* homologs *mshDOP*. *mshDOPQ* exhibit conserved gene synteny with T4aP systems encoding *pilVWXY1* in the *Pseudomonadales* order, indicating potential functional homology between these systems (**Fig. S1**). AlphaFold modeling predicts that the extreme C-terminus of MshQ, like PilC/PilY1, engages in β-strand complementation with the PilK homolog MshP (**Fig. S10**). This prediction is in accord with β-strand complementation regulating tip complex assembly in divergent T4P, and suggests that AlphaFold modeling of tip complexes may enable identification of as-of-yet unidentified tip complex subunits.

*Bacillus subtilis* Com T4dP, and *V. cholerae* T4aP, both involved in DNA uptake, do not include a PilC/PilY1 ortholog. In each case the PilK ortholog has an additional β-strand, similar to T2SS GspK pilins. One of the most divergent subgroups of T4F include the Tad T4cP, which until recently were thought to contain only two minor pilin tip complex subunits (43). The minor pilin-like protein CpaL (∼600 residues) was recently identified as a component of the *C. crescentus* Tad T4cP tip complex, playing an important role in bacterial surface sensing (43). Because Tad T4cP represent the most divergent evolutionary clade of bacterial T4P, we reasoned that the presence of C-terminal β-strand complementation in this clade might suggest β-strand complementation as a mechanism present in the ancestral T4F system (**Fig. S9**) (4). As in other T4F systems, AlphaFold modeling of CpaL with previously identified minor pilin subunits CpaJ and CpaK predicts that the extreme C-terminus of CpaL exhibits β-strand complementation with the β-sheet in its N-terminally located minor pilin-like domain (**Fig. S10, S15, 5A**). Additional AlphaFold modeling suggests that the middle ∼470 residues of CpaL may be dispensable for tip complex stability, and predicts that a fusion of the extreme C-terminal six amino acids to the minor pilin-like domain might function like T2SS GspK or like our chimeric PilK/X-Cβ mutants. Using a hyperpiliated mutant of *C. crescentus* (44), we constructed Δ*cpaL* and *cpaL*Δ*Cβ* strains, as well as a strain lacking the middle region of CpaL at the native locus (Δmiddle) (**Fig. 5**). Similar to T4aP systems lacking the C-terminal β-strand, T4P were not detected in the Δ*cpaL* strain and were almost undetectable in the *cpaL*Δ*Cβ* mutant. In contrast, the Δmiddle strain produced significantly more T4P, with piliation restored in almost 15% of the population (∼38% of wild-type levels).

**Figure 5.**
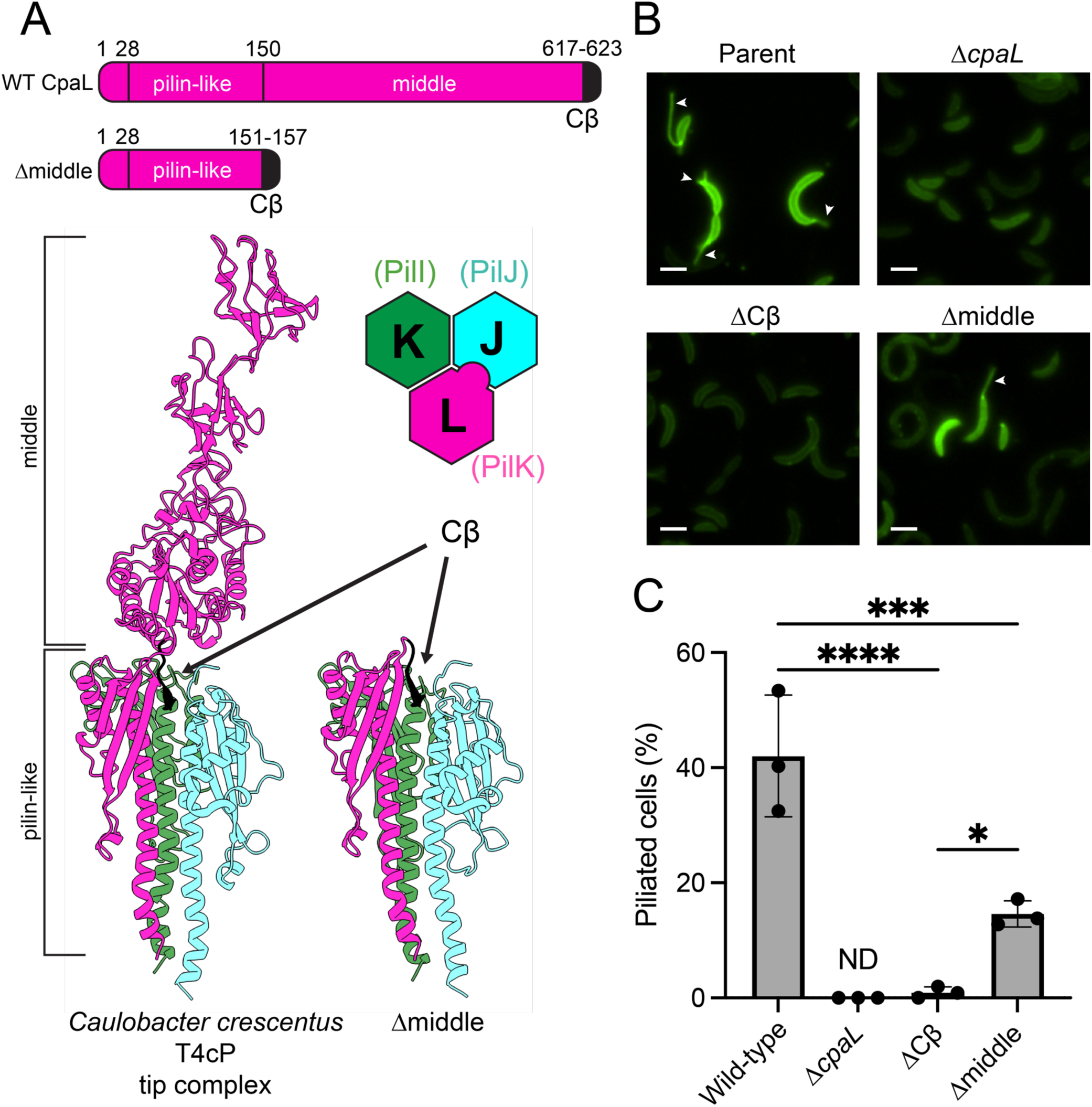
Diverse T4Fs employ C-terminal β-strand complementation to stabilize the tip complex. (A) A schematic of wildtype CpaL (WT) compared to the Δmiddle strain lacking the internal ∼470 residues predicted to sit atop the T4cP tip (top). AlphaFold models of the wildtype or Δmiddle tip complexes from *C. crescentus* CB13. In both models, the conserved C-terminal β-strand is colored black. (B) Representative microscopy images of indicated strains with background fluorescence subtracted. White arrows indicate examples of visible T4P filaments. Scale bars, 2 µm. (C) Quantification of piliated cells in populations of strains shown in (B). Bar graph shows the mean +/- SD. Each dot represents an independent, biological replicate. Statistics were determined by one-way ANOVA and corrected by Sidak’s multiple comparisons test. ****P<0.0001; ***P<0.001, *P<0.05; ND, not detected.

## DISCUSSION

Our results indicate that C-terminal β-strand complementation within diverse T4F tip complexes is important for initiating fiber polymerization and might have been a feature of ancestral T4F systems. The T4F superfamily is among the most versatile and broadly distributed groups of nanomachines found in prokaryotes and was likely present in the last universal common ancestor of life. However, it is still unclear how these dynamic structures are regulated and what signals trigger initiation of fiber polymerization. In this work we uncovered a crucial aspect of fiber assembly by identifying β-strand complementation as a key mechanism of tip complex licensing and stability. We show β-strand complementation can be provided by either a tip-associated functional protein like those in the PilC/PilY1 family or by the most apical minor pilin (or minor pilin-like protein) in the complex, as is the case for T2SS and Tad T4cP. The conservation of β-strand complementation as a mechanism for promoting tip complex assembly between the distantly related Tad T4cP and other T4F suggests that ancestral T4F from which all modern T4Fs descended may have used a similar mechanism to promote fiber assembly (**Fig. S9**). Although not analyzed in the present study, most Pil/Gsp operons that encode I, J, and K subunits also encode an H subunit. In T2SS tip complexes, GspH sits at the GspJ-GspK interface (32, 45). We speculate that in T4P, PilH also binds at the J/K interface, and thereby monitors the assembly status of the quaternary PilC/Y1-PilK/PilI/PilJ complex.

Incomplete rescue of fiber initiation by PilK/X-Cβ chimeras suggests that PilC/PilY1 proteins have additional functions in T4P biogenesis and dynamics, and we suggest that the conserved PilC/PilY1 β-propellor domain accelerates or stabilizes formation of the tip initiation holocomplex. Like the conserved C-terminal β-propellor of PilC/Y1, the platform of some T2SS substrates, including cholera toxin and *E. coli* heat labile toxin, have β-propellor folds. However, these secreted proteins do not require strand complementation on PilK for association with the tip complex. This is consistent both with a conserved role for the β-propellor in fiber initiation or extension, and with the requirement for T2SS payloads to be released from the extended pseudopilus rather than stably retained at the fiber tip. The β-propellor may accelerate or stabilize assembly of the tip complex, promote gating of the secretin pore (20), or both. However, neither the β-propellor nor the C-terminal β-strand are formally essential for fiber initiation, since strains lacking PilC/PilY1 or core minor pilins, are piliated when retraction is prevented by removal of the PilT retraction ATPase. The idea that PilC/PilY1 promote assembly of minor pilins in the tip complex is strongly supported by observations that pili which assemble in the absence of both PilC/PilY1 and the retraction ATPase PilT are depleted of tip-associated minor pilins (17).

Bioinformatic analyses of T4F systems have provided a roadmap of evolutionary trajectories for diverse T4F (4). AlphaFold modeling provides a remarkable platform for predicting how tip complexes have diverged over time and across ecological niches. Our models of diverse T4F suggest the MSHA protein MshQ may act as a functional homolog of PilC/PilY1 proteins (**Fig. S10**). Viewed in the context of T4F phylogenies, these models suggest that PilC/PilY1 and other large adhesins evolved after the divergence of Com T4dP from the ancestor of modern T4aP/T2SS/MSHA pili. However, the absence of a functional homolog in T4aP required for DNA uptake from *V. cholerae*, and in T2SS which fall within this clade, may suggest that tip adhesins were subsequently lost from certain lineages (**Fig. S9**).

In solved structures of T4bP tip complexes from the *V. cholereae* toxin <u>c</u>o-regulated pilus (TCP) and the <u>c</u>olonization factor <u>a</u>ntigen/III (CFA/III) pilus found in human enterotoxigenic *E. coli*, the filament tip is composed of a minor pilin homotrimer with fundamentally distinct pilin-pilin interactions (46). These T4P gene loci only contain a single minor pilin gene, in contrast with most T4F systems which harbor multiple minor pilins, encoded within syntenic operons. But some T4bP operons do encode PilIJK homologs, including the <u>b</u>undle-forming <u>p</u>ili (BFP) found in some *E. coli* lineages. Together, these observations suggest that the ancestral T4bP diverged from other T4F tip complex assembly mechanisms early in evolution, consistent with phylogeny-based predictions (**Fig. S9**) (4). This interpretation is strengthened when we examine T4cP and T4dP.

T4dP Com (competence) pili are unique to Gram-positive bacteria. For *Streptococcus* spp., modeling and experiments support the formation of a tip complex containing PilIJK homologs ComGE/GF/GG (6, 45). ComGF is needed for DNA binding (45, 47, 48). Similarly, the *B. subtilis* Com operon comprises ComGE/GF/GG proteins. As in *S. pneumoniae*, *B. subtilis* ComGF contains a canonical prepilin signal sequence. Our AlphaFold modeling confidently predicts the formation of a *B. subtilis* tip complex. However, processed *B. subtilis* ComGF lacks the typical pilin N-terminal domain. Our *B. subtilis* model (**Fig. S10**) confidently predicts that the extreme C-terminal β-strand of ComGG behaves similarly to the T2SS GspK homolog, Curiously, however, the *S. pneumoniae* ComGG was previously modeled with low confidence as an α-helix, not a β-strand (6, 45). Nevertheless, ComGF interacts with GE and GG in the same orientation in models of both *Streptococcus* and *B. subtilis* tip complexes. Thus, it appears likely that β-strand complementation occurs in at least part of the Com T4dP clade (**Fig. S9, S10**). Similarly, our modeling and experiments show that the minor pilin-like protein CpaL from Tad T4cP systems acts as a PilK analog (**Fig. S1, Fig. 5**)(51). In summary, β-strand complementation in PilK paralogs may occur, *in cis* or *in trans*, in T2SS, T4aP, T4bP, T4cP, and T4dP. We therefore suggest that licensing mechanisms like those characterized here are common to many or even most T4F systems, and that these mechanisms arose early in the evolution of T4F.

## MATERIALS & METHODS

### Bacterial strains and culture conditions

*Neisseria gonorrhoeae* strain FA1090, *Acinetobacter baylyi* strain ADP1, and the hyperpiliated derivative *Caulobacter crescentus* CB13 strain bNY30a were used in this study. For a list of strains and primers used throughout, see **Tables S1 and S2** respectively. *N. gonorrhoeae* was grown on GCB agar with Kellogg’s supplements (47). *A. baylyi* cultures were grown at 30 °C in Miller lysogeny broth (LB) medium and on agar supplemented with kanamycin (50 µg/ml), spectinomycin (60 µg/ml), zeocin (50 µg/ml), and apramycin (50 µg/ml) where appropriate. *C. crescentus* was grown in peptone yeast extract (PYE) medium and on agar supplemented with kanamycin (5-10 µg/ml) and nalidixic acid (20 µg/ml) where appropriate.

### Strain construction

#### N. gonorrhoeae

Mutations were introduced in the FA1090 background by spot transformations of PCR products and/or linearized plasmids. Transformed spots were streaked to GCB plates or GCB plates with the appropriate antibiotic then colonies were screened by sequencing of the mutated locus.

To construct the pilE-T125C strains AMN64 and AMN65, pilE(1-81-S2) was amplified and mutated to T125C by overlap extension PCR. To disrupt pilT in the pilE-T125C background, the ΔpilT::ermC allele was amplified directly from AMN4 and used to transform AMN65. All other mutations were introduced by transformation of plasmids constructed by Gibson assembly of PCR products, gBlocks (IDT) and/or extended oligonucleotides into pBluescript KS(+).

For transformation assays pAMP2097was prepared. This plasmid contains the gyrB-D429N allele, which confers resistance to nalidixic acid, and a *Neisseria* DNA uptake sequence. *N. gonorrhoeae* AMN18 genomic DNA was used as template for PCR with primers MS11_gyrB-F and MS11_gyrB-R. The PCR product was purified, digested with EcoRI and ClaI (NEB) and cloned into pBluescriptKS+ (Stratagene), yielding AMP2096. Site-directed mutagenesis of AMP2096 with primers gyrB_D429N-f and gyrB_D429N-r was used to introduce the D429N mutation, resulting in pAMP2097.

#### A. baylyi

Mutants in *A. baylyi* were made using natural transformation as described previously (48). Briefly, mutant constructs were made by splicing-by-overlap (SOE) PCR to stich (1) ∼3 kb of the homologous region upstream of the gene of interest, (2) the mutation where appropriate (for deletion by allelic replacement with an AbR cassette), and (3) ∼3 kb of the homologous downstream region. The upstream region was amplified using F1 + R1 primers, and the downstream region was amplified using F2 + R2 primers. All AbR cassettes were amplified with ABD123 (ATTCCGGGGATCCGTCGAC) and ABD124 (TGTAGGCTGGAGCTGCTTC). In-frame deletions were constructed using F1 + R1 primer pairs to amplify the upstream region and F2 + R2 primer pairs to amplify the downstream region with ∼20 bp homology to the remaining region of the downstream region built into the R1 primer and ∼20 bp homology to the upstream region built into the F2 primer. SOE PCR reactions were performed using a mixture of the upstream and downstream regions, and middle region where appropriate using F1 + R2 primers. SOE PCR products were added with 50 µL of overnight-grown culture to 450 µL of LB in 2-mL round-bottom microcentrifuge tubes (USA Scientific) and grown at 30 °C rotating on a roller drum for 3–5 h. For AbR-constructs, transformants were serially diluted and plated on LB and LB + antibiotic. For protein fusion constructs, after the 3–5 h incubation, cells were diluted and 100 µL of 10^−6^ dilution was plated on LB plates. Deletions were confirmed by PCR using primers ∼150 bp up- and downstream of the introduced mutation.

#### C. crescentus

In-frame deletion strains were constructed by double homologous recombination using pNPTS-derived plasmids as previously described (44). S-17 *E. coli* cells harboring pNPTS-derived plasmids were grown to exponential phase and then 100 µl was pelleted by centrifugation at 18,000*g* for 1 min. One ml of overnight culture of *C. crescentus* was pelleted by centrifugation at 5,200*g* for 1 min, and then both pellets were resuspended in the same 10 µl of fresh PYE and spotted onto a PYE 1.5% agarose plate. Plates were incubated overnight at 30 °C, and the next day cells were removed from the plate and resuspended in 100 µl PYE before plating on PYE plates containing kanamycin and nalidixic acid. After three days of growth at 30 °C, colonies were patched onto fresh kan/nal PYE plates and grown overnight at 30 °C. The next day, cells from each patched colony were inoculated into 3 ml PYE medium and grown overnight at 30 °C. 10 µl of overnight culture was then spotted onto PYE plates containing 3% sucrose and struck out for single colony isolation. Sucrose plates were grown for three days at 30 °C and then 4-6 colonies from each plate were patched onto PYE plates +/- kanamycin and incubated overnight at 30 °C. Colonies that were kanamycin sensitive were confirmed by PCR and nanopore sequencing

For construction of the pNPTS-derived plasmids, ∼500 bp flanking regions of DNA on either side of the desired mutations were amplified from bNY30a DNA. Upstream regions were amplified using F1 and R1 primers while downstream regions were amplified using F2 and R2 primers. The resulting amplified DNA was purified (Qiaquick) and assembled into pNPTS138 that had been digested with restriction enzyme *Eco*RV (New England Biolabs) using HiFi Assembly Master Mix (New England Biolabs).

### Modeling

Preliminary modeling was done using AlphaFold2 in the CollabFold environment (49). The modeling shown in the figures was done using AlphaFold3 (50) on the DeepMind server. Protein processing sites were predicted through a combination of manual analysis, freely available prediction software SignalP 6.0 (51), and/or DeepTMHMM (52). All protein sequences used for AlphaFold models presented in this work can be found in **Extended Dataset 1**. All models were generated for display using ChimeraX software (53).

### Minor pilin expression in *E. coli*

Minor pilin proteins were produced in soluble form with oxidized disulfide bonds in the *E. coli* periplasm as described (34). Size exclusion chromatography was done on a calibrated Superdex 75 Increase 10/300 column (Cytiva). For co-expression of soluble PilK-His_6_ and PilC, PilK-His_6_ was expressed from a plasmid previously described (34), and FLAG-PilC(533-1060) bearing an N-terminal signal peptide was expressed from a low-copy plasmid [DETAILS]

### Live cell imaging and in vivo labeling of dynamic T4P

#### N. gonorrhoeae

Pilus labeling was as described (21), with modifications. Briefly, *N. gonorrhoeae* cells expressing Cys-substituted PilE major pilin were grown on GCB agar with Kellogg supplements and resuspended into Labeling Medium: DMEM (FluoroBrite, Gibco A18967-01) that had been supplemented with 1× GlutaMAX (Gibco 35050-061) and 30 mM HEPES, and pre-equilibrated to 37°C and 5% CO_2_. The cell suspension was adjusted to OD_600nm_ ∼0.2 in 1mL, in 1.8 mL round-bottom microcentrifuge tubes. The cells were sedimented (7,000rpm, 1 min.) and resuspended in Labeling Medium containing 20 µM Alexa Fluor 488 C5 Maleimide (Thermo-Fisher #A10254, FW=721), from a 5 mM stock solution prepared in anhydrous DMSO. The cells were incubated at 37°C and 5% CO_2_ with periodic gentle swirling of the cells for 15min. The labeled cells were sedimented (7,000 rpm, 1 min.), washed in 1 mL of Labeling Medium, again sedimented, and washed for a total of three times before finally being resuspended in 200 µL of Imaging Medium: Labeling Medium, further supplemented with 500 µM TEMPOL (Enzo #ALX-431-081G001), 1 mM Trolox (Vector Labs CB-01000-2), 500 µM freshly prepared ascorbic acid, and 0.3% bovine serum albumin. For some samples, wells were additionally pre-treated with 0.01% poly-L-lysine to stick bacteria to the coverslip. T4P were observed using a Yokogawa CSU-X1 spinning disk confocal head, with a 100/1.45 NA Plan Apochromat objective. For excitation a 488 nm diode laser was coupled to the confocal head using a single mode fiber. The incident illumination power was calibrated at the objective back aperture using a laser power meter (Thorlabs) and corresponded to 15-20 W/cm^2^ at the sample plane (54). Image sequences were captured using an Andor 888 EMCCD camera operated at 25 frames/s. The microscope system was controlled, and data were acquired, using NIS Elements software.

#### A. baylyi

Pilin labeling in *A. baylyi* was performed as described previously (39). Briefly, 100 µL of overnight cultures was added to 900 µL of fresh LB in a 1.5 mL microcentrifuge tube, and cells were grown at 30 °C rotating on a roller drum for 70 min. Cells were then centrifuged at 18,000*g* for 1 min and resuspended in 50 µL of LB before labeling with 25 µg/mL of AlexaFluor488 C5-maleimide (AF488-mal) (ThermoFisher) for 15 min at room temperature. Labeled cells were centrifuged, washed three times with 100 µL of PBS and resuspended in 5–20 µL PBS. Cell bodies were imaged using phase-contrast microscopy while labeled pili were imaged using fluorescence microscopy on a Nikon Ti2-E microscope using a Plan Apo 100X oil immersion objective, a GFP/FITC/Cy2 filter set for pili, a Hamamatsu ORCA-Fusion Gen-III cCMOS camera, and Nikon NIS Elements Imaging Software. Cell numbers and the percent of cells making pili were quantified manually using Fiji. All imaging was performed under 1% agarose pads made with PBS solution.

#### C. crescentus

Briefly, 200 μl of overnight cultures was added to 3 mL of fresh PYE medium in a 14 mL culture tube, and cells were grown at 30 °C rotating on a roller drum for 3 h. 1.5 mL of each culture was then centrifuged at 5,200 x *g* for 1 min and resuspended in 50 μl of PYE before labeling with 25 μg/ml AlexaFluor488 C5-maleimide (AF488-mal) (ThermoFisher) for 30 min at room temperature. Labeled cells were centrifuged, washed one time with 200 μl of PYE, and resuspended in 10–40 μl of PYE. Cell bodies were imaged employing the same microscopy setup as described above. Cell numbers and the percentage of cells making pili were quantified manually using Fiji. All imaging was performed under 1% agarose pads made with water.

The percentage of cells expressing T4P for all species was manually quantified using Fiji software (55). Micrograph images of T4P were analyzed using Nikon NIS Elements software and ImageJ. Micrographs are representative of each strain and are presented with intensity equalized over time, as well as a rolling median over every 3 frames to minimize background noise. The LUTs for *N. gonorrhoeae* strains were adjusted for each strain to optimize visualization of T4P present. For *A. baylyi* and *C. crescentus*, LUTs were normalized for each species.

### Immunofluorescence in *ΔfimL ΔpilT* mutants

We previously found that overnight cultures of *A. baylyi* grown in M63 minimal medium + casamino acids and glucose (M63CA + G) (2 g/l (NH_4_)_2_SO_4_, 13.6 g/l KH_2_PO_4_, 0.5 mg/l FeSO_4_, 1 mM MgSO_4_, 0.5% w/v casamino acids, and 0.4% w/v glucose) produce longer pili than cells grown in LB (39). We therefore grew cells in M63CA + G medium for assessing PilY1 localization on *A. baylyi* pili as this increase in length vastly improved our ability to resolve individual pilus fibers. For immunofluorescence experiments, 100 μl of overnight cultures grown in M63CA + G was centrifuged and resuspended in 100 μl of PBST (PBS + 0.05% v/v Tween-20) containing a 1:100 dilution of mouse monoclonal α-FLAG antibodies (Sigma) and incubated overnight at 4 °C with end-over-end rotation in a microcentrifuge tube. The next day, cells were washed three times with 200 μl of cold PBST before resuspension in 100 μl of cold PBST containing a 1:200 dilution of goat α-mouse antibodies conjugated to AlexaFluor555 (Abcam). Cells were incubated for 2 h at 4°C with end-over-end rotation before washing three times with 200 μl of cold PBST. Cells were then resuspended into 50 μl of cold PBST and incubated with 25 μg/ml AF488-mal for 15 min. Cells were then washed once with plain PBS (Tween-20 was omitted here as it has considerable background fluorescence) and then resuspended in 5–30 μl of plain PBS and imaged. A 1 μl aliquot of cells was used for microscopy employing the same microscopy setup as described above, using a DsRed/TRITC/Cy3 filter set (Nikon).

### Transformation assays

#### N. gonorrhoeae

Cells were grown on GCB agar with Kellogg supplements and resuspended into 500 µL GCBL media supplemented with 5 mM MgSO_4_. Cell suspensions were adjusted to 1.2×10^7^ cfu/ml. Suspensions were then incubated with 250ng linearized AMP2097 for 15min at 37° C 5% CO_2_ in a 96-well plate format (Falcon #351172). The transformation mixture was then diluted 1:10 into GCBL media supplemented with Kellogg supplements and incubated for 4 hours at 37 °C 5% CO_2_. Subsequent 1:10 serial dilutions of the transformation reaction mix were performed into GCBL media supplemented with Kellogg supplements. The serial dilutions were plated using the “dribble plating” method described previously onto 2 sets of plates: GCB plates and plates containing GCB supplemented with 1.5 µg/ml Nalidixic acid (both supplemented with Kellogg supplements). Plates were left to grow for two nights at 37°C 5% CO_2_ before assessing colony growth. For quantitative transformation in the presence of IPTG, cells were grown on GCB agar supplemented with 1 mM IPTG on the surface of the agar. Subsequent GCBL media used for resuspension, transformation, and serial dilutions were also supplemented with 1 mM IPTG. Plates used for dribble plating did not contain IPTG.

#### A. baylyi

Transformation assays in *A. baylyi* were performed exactly as previously described (39, 48). Briefly, strains were grown overnight in LB broth at 30°C on a roller drum. Then, 50 μl of overnight culture was subcultured into 450 μl of fresh LB medium, and at least 50 ng of transforming DNA (tDNA) (an ∼7 kb PCR product containing Δ*pilT*::*spec* amplified using primers CE49 + CE50) was used for all transformation assays except for DNA competition assays which are detailed below. DNA was quantified using a Qubit (ThermoFisher) following standard Qubit protocols. Reactions were incubated with end-over-end rotation on a roller drum at 30°C for 5 h and then plated for quantitative culture on LB + antibiotic plates (to quantify transformants) and on plain LB plates (to quantify total viable counts). Data are reported as the transformation frequency, which is defined as the [colony-forming units (CFU/ml of transformants)/(CFU/ml of total viable counts).

### Western blotting in *A. baylyi*

50 µl of overnight cultures were diluted into 3 mL fresh LB media and grown to exponential phase at 30 °C for 3 hours. The cultures were then normalized to an OD_600_ of 1.0 in a volume of 500 µl and concentrated into a pellet by centrifugation, and the culture supernatant was discarded. Cell pellets were resuspended in 50 µl PBS and then mixed with an equal volume of SDS-PAGE sample buffer (125 mM Tris, pH 6.8, 20% glycerol, 4% SDS, 0.4% bromophenol blue, and 10% β-mercaptoethanol) and denatured using a heat block set to 99 °C for 10 min. Proteins were separated on a 4–20% pre-cast polyacrylamide gel (Biorad) by SDS electrophoresis, electrophoretically transferred to a nitrocellulose membrane, and probed with 1:5000 dilution of mouse monoclonal α-FLAG antibodies (Sigma) and a 1:12000 dilution of mouse monoclonal α-RpoA (BioLegend) primary antibodies. Blots were washed and then incubated in a 1:10000 dilution of goat α-mouse antibody conjugated to horseradish peroxidase secondary antibody (Sigma). Blots were washed again and then incubated with SuperSignal West Pico PLUS Chemiluminescence substrate (ThermoFisher). Blots were then imaged using a BioRad Chemidoc imaging system.

### Statistics

All statistical analyses were performed using Prism software version 10.4.1.

## ACKNOWLEDGEMENTS

We thank K. Forest, H. Seifert, and R. Klevit for helpful discussions, D. Miller for suggesting the term “licensing,” and M. So and H. Seifert for sharing materials. This work was funded by R21 AI155991 (AJM), R35 GM150916 (CKE), NIH R01AI150152 (BT), T32AI07410 (JA), T32GM153507 (ZRL), University of Washington School of Medicine Molecular Biophysics Training Program Hurd Fellowship (ZRL), University of Washington School of Medicine Medical Scientist Training Program (T32 GM007266 and T32 GM153182) and the University of Washington Bacterial Research Fund, including a gift from Dr. J.M. Griffiss (SM, AJM). Courtney Ellison, PhD, is a Damon Runyon-Marilyn and Scott Urdang Breakthrough Scientist supported by the Damon Runyon Cancer Research Foundation (DFS6023)

---

**Figure S1.**
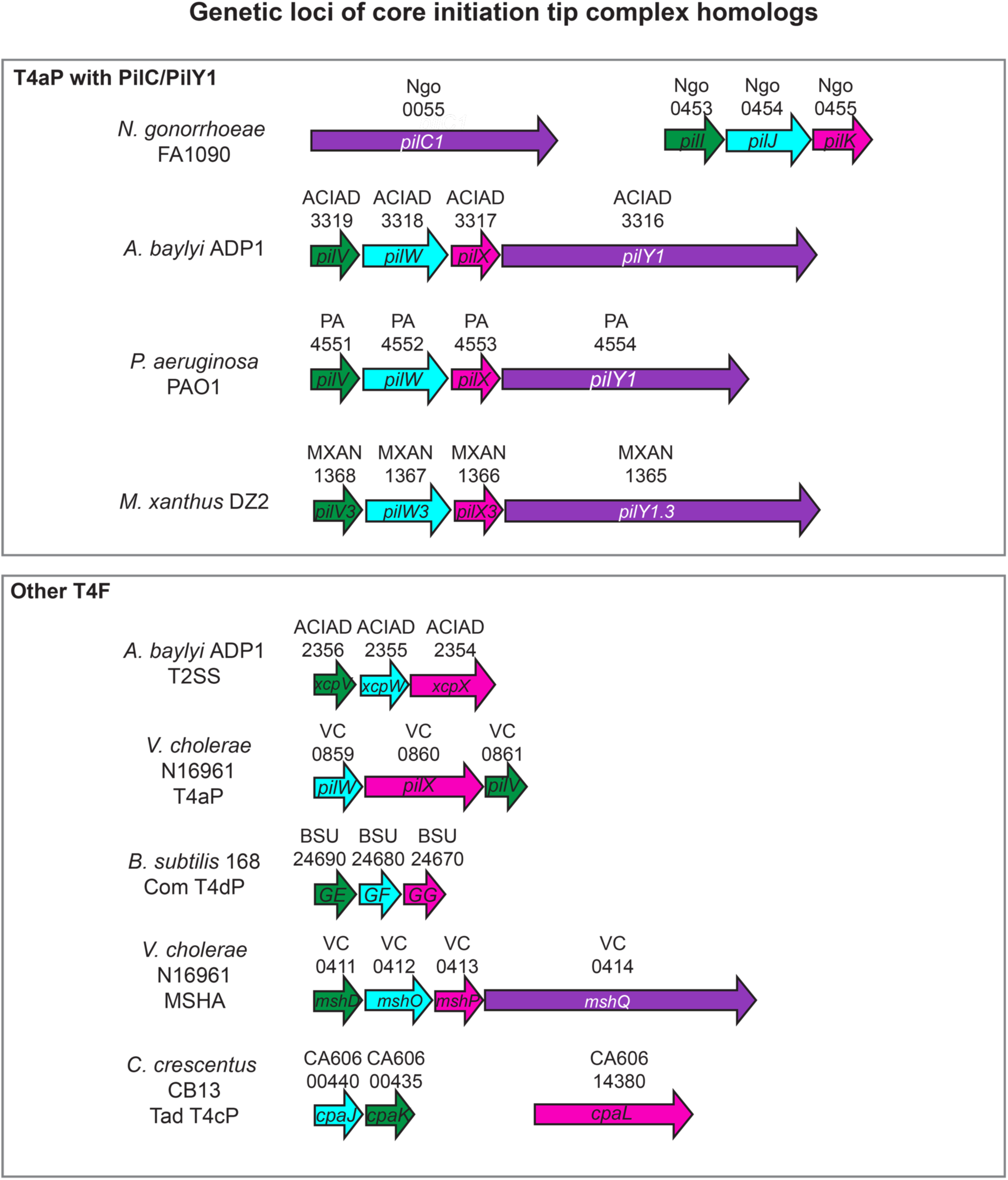
Schematic of genetic loci for several T4F discussed in this manuscript. Numbers above genes indicate locus tags. Protein homologs are colored correspondingly.

**Figure S2.**
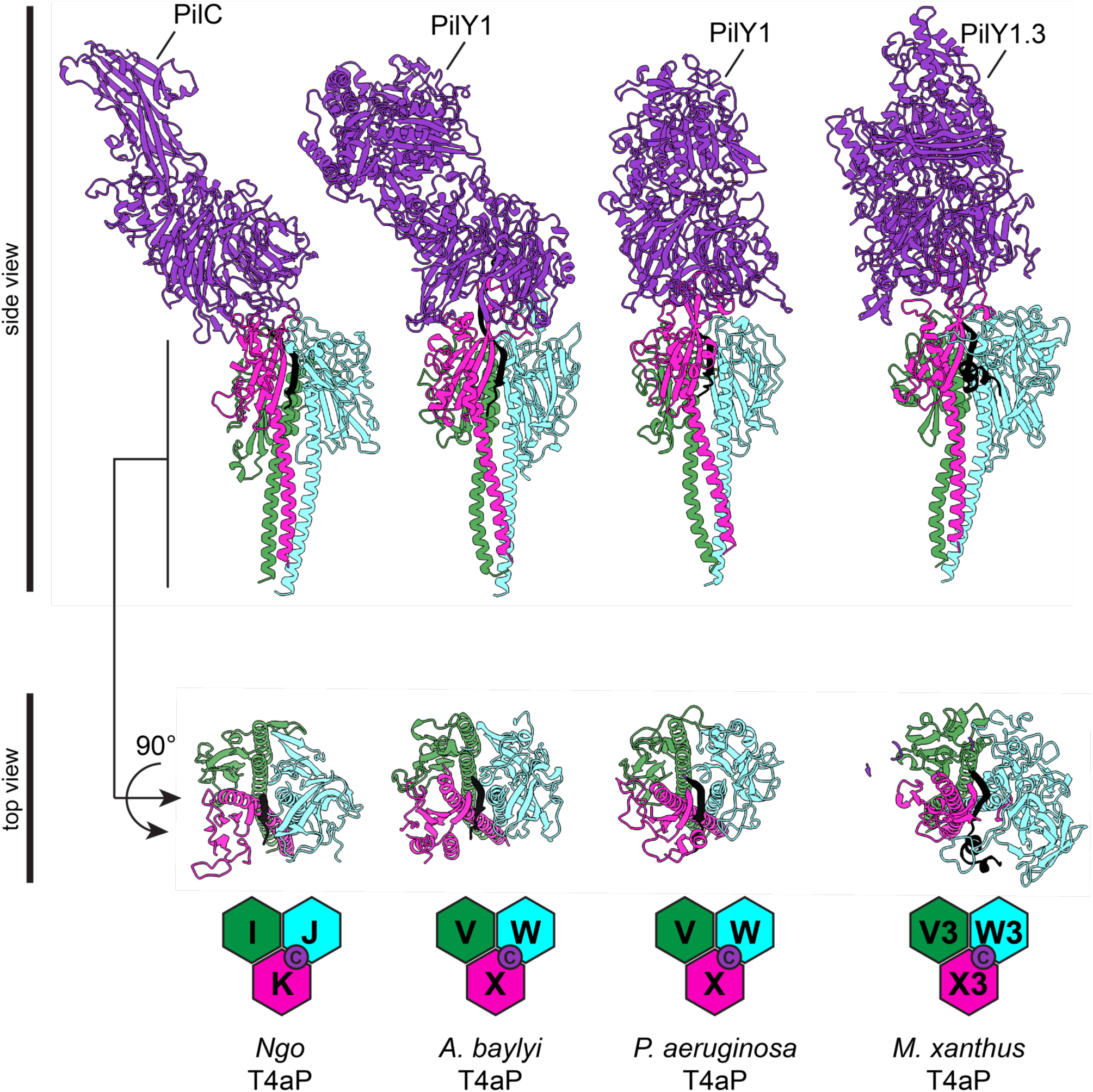
A conserved β-strand is predicted to strand-complement the β-sheet in GspK homologs of T4aP. AlphaFold-predicted models of minor pilins in complex with PilC or PilY1. In all models, the conserved C-terminal β-strand is colored black.

**Figure S3.**
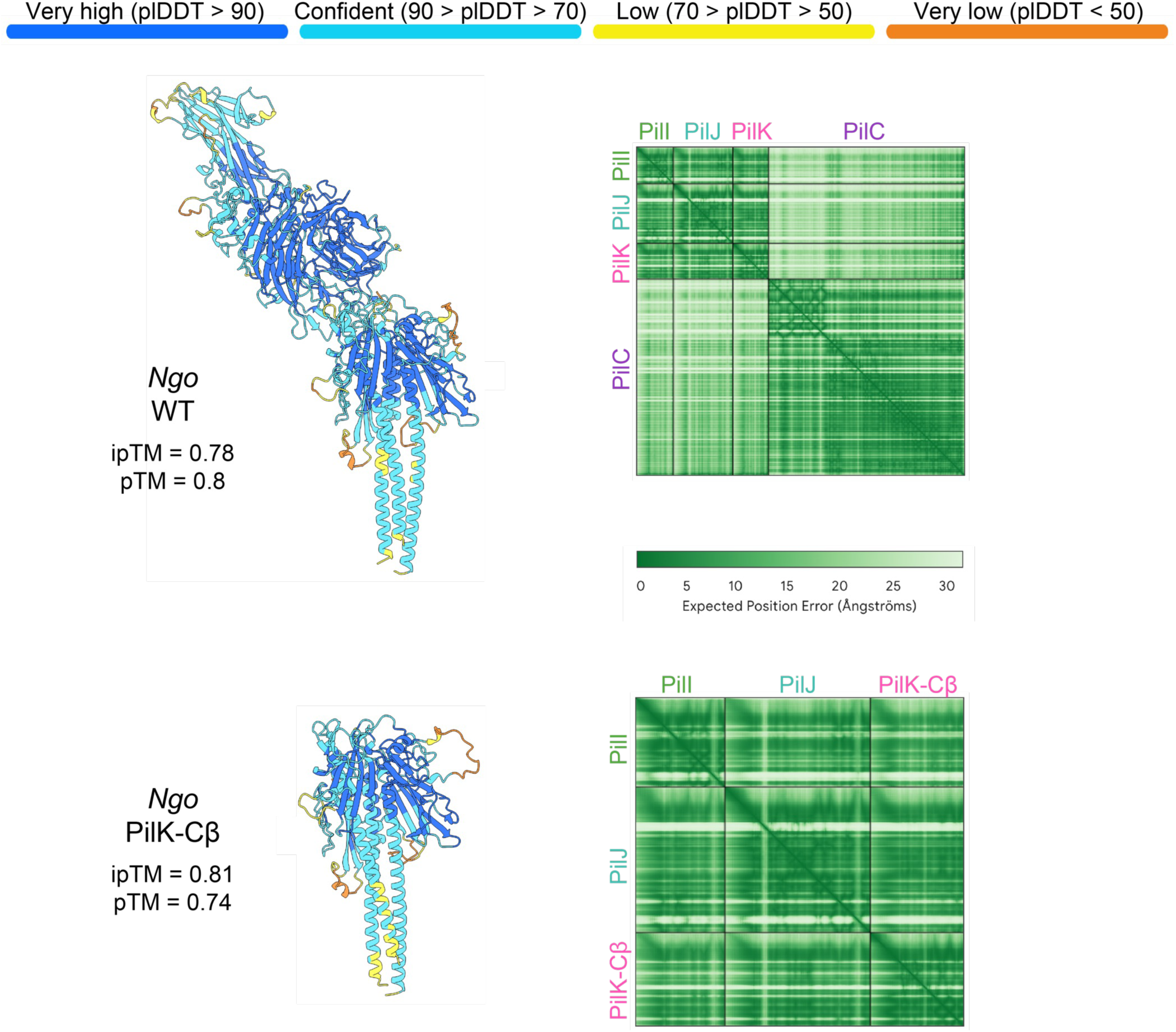
AlphaFold confidence metrics for tip complexes from wildtype T4aP in *N. gonorrhoeae* and the PilK-Cβ mutant used in this work. Structures on the left are colored by confidence intervals indicated at the top. The boxes on the right show predicted aligned error of structures on the left.

**Figure S4.**
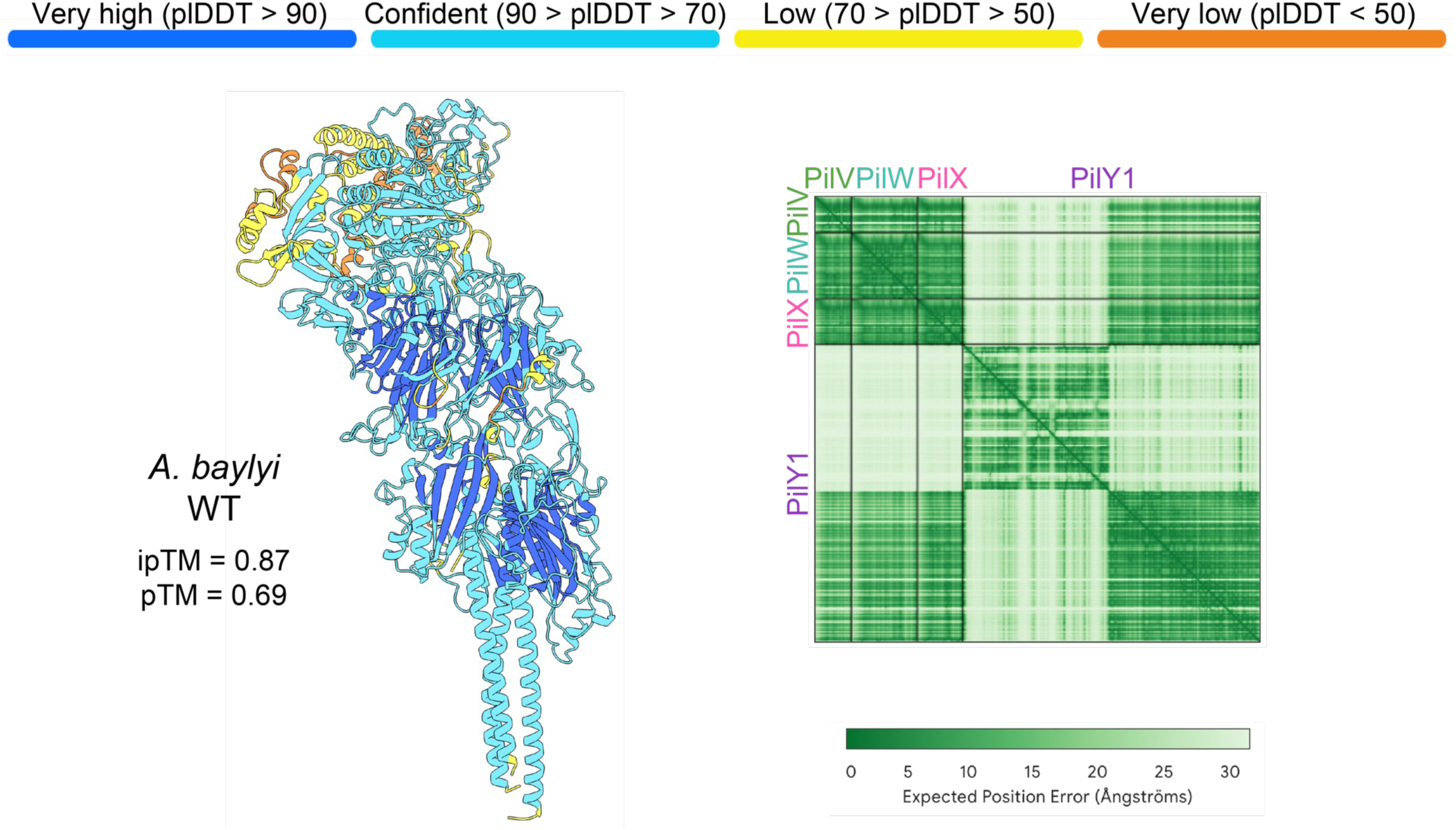
AlphaFold confidence metrics for the tip complex from wildtype T4aP in *A. baylyi.* Structure on the left is colored by confidence intervals indicated at the top. The box on the right shows the predicted aligned error of structure on the left.

**Figure S5.**
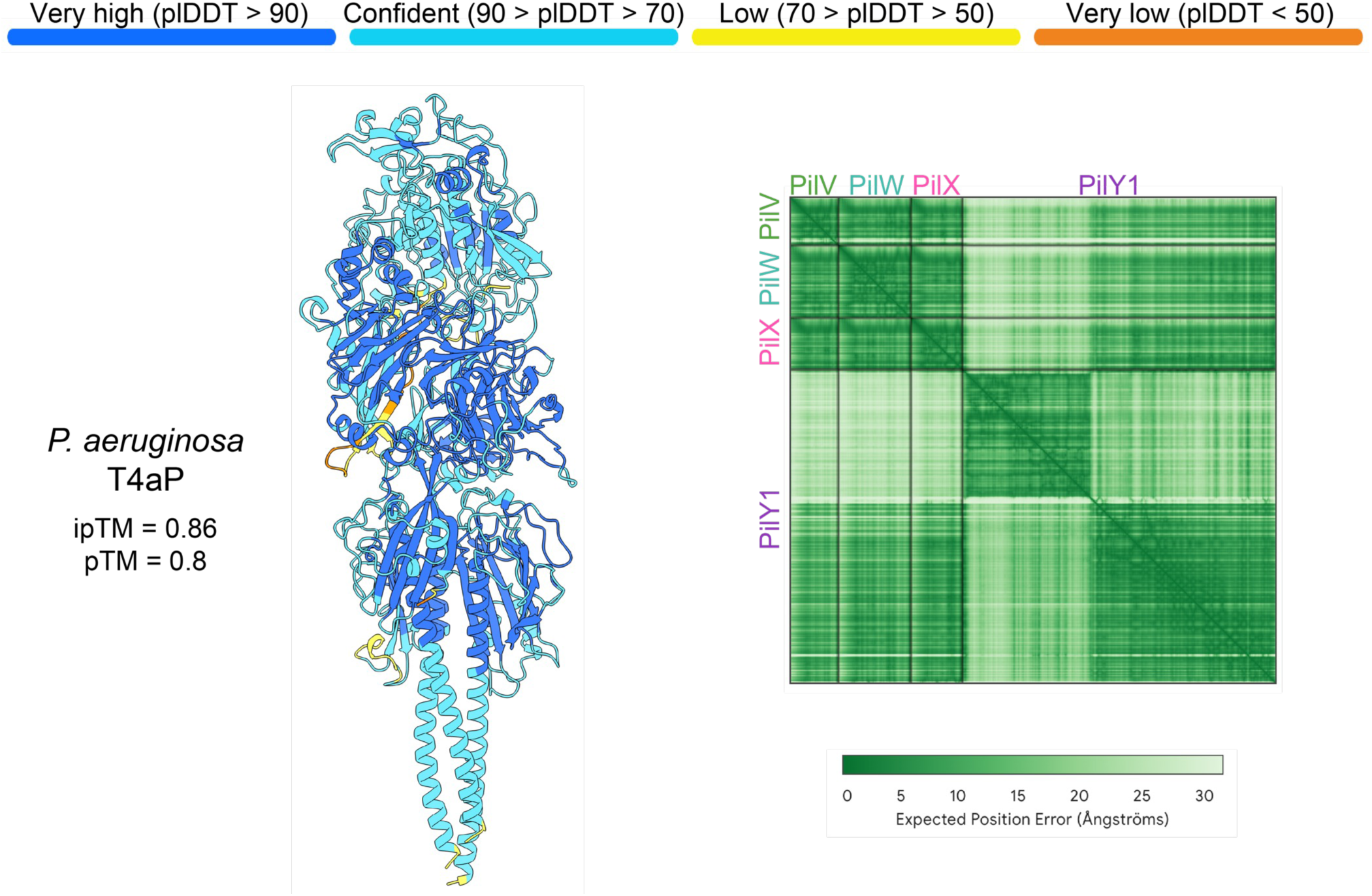
AlphaFold confidence metrics for the tip complex from wildtype T4aP in *P. aeruginosa* Structure on the left is colored by confidence intervals indicated at the top. The box on the right shows the predicted aligned error of structure on the left.

**Figure S6.**
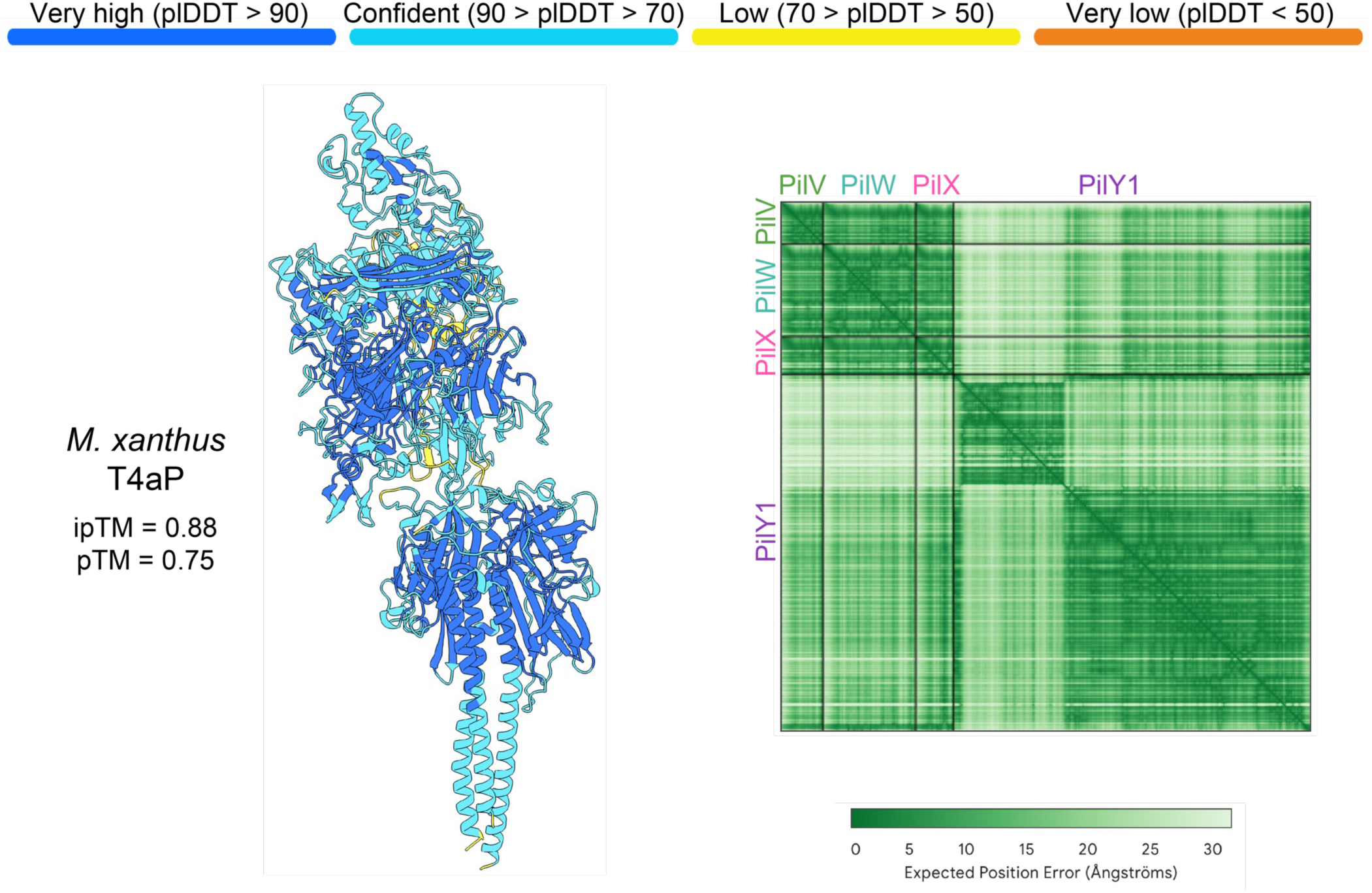
AlphaFold confidence metrics for the tip complex from wildtype T4aP in *M. xanthus*. Structure on the left is colored by confidence intervals indicated at the top. The box on the right shows the predicted aligned error of structure on the left.

**Figure S7.**
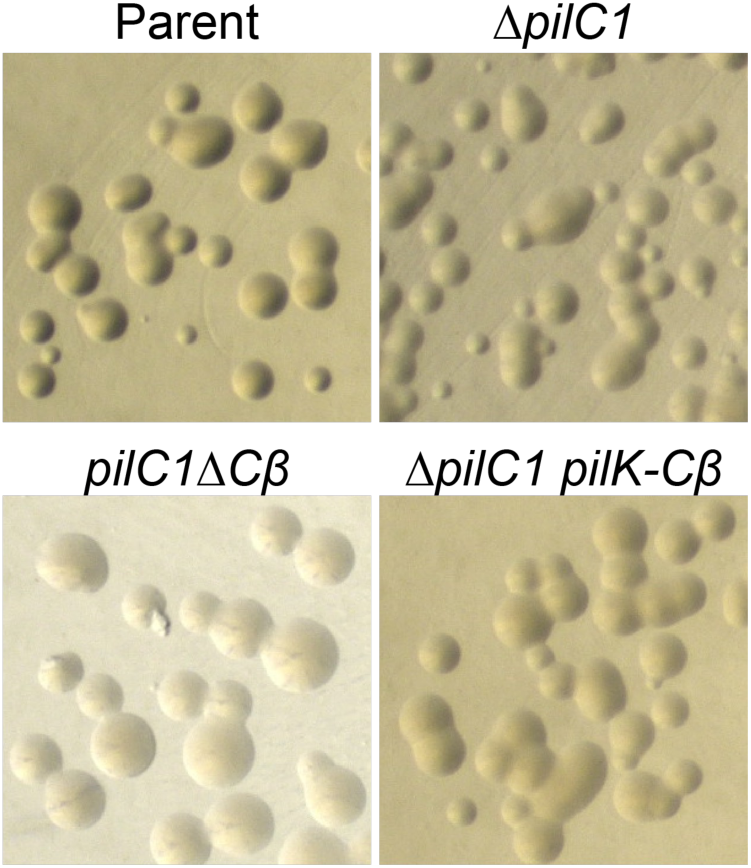
Colony morphology of *N. gonorrhoeae pilC* mutants and the PilK-Cβ strain.

**Figure S8.**
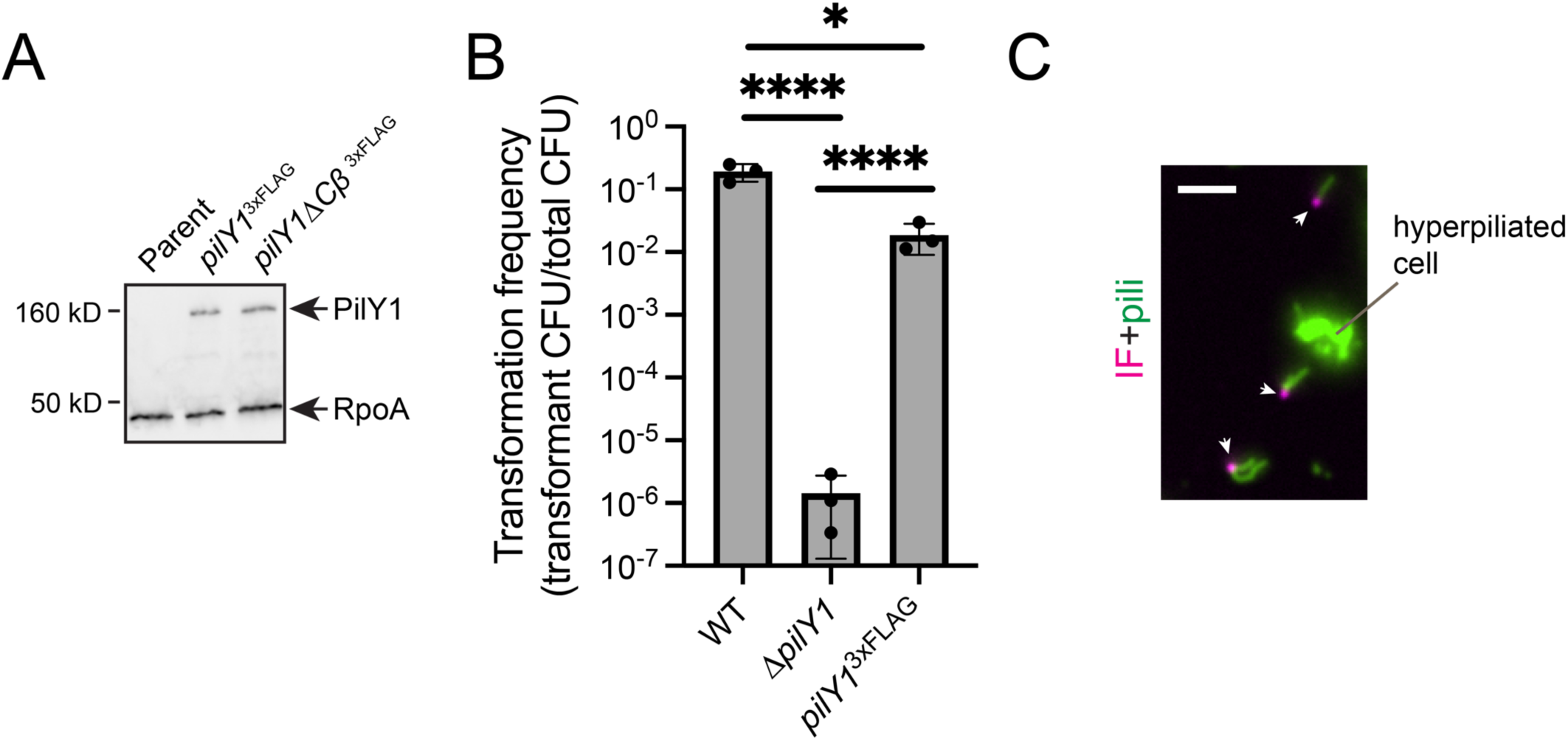
PilY1^3xFLAG^ and immunofluorescence supplementary data. (A) Representative Western analysis of indicated strains. RpoA was used as a loading control. Blot was probed with both ⍺-FLAG and ⍺-RpoA antibodies. (B) Natural transformation assay data from indicated strains. Bar graph shows the mean +/- SD. Each dot represents an independent, biological replicate. Statistical analysis for natural transformation assays was performed on log-transformed transformation frequency data. Statistics were determined by Sidak’s multiple comparisons test. ****P<0.0001; *P<0.05. (B) Representative microscopy image of sheared T4P with tip-associated PilY1 signal. White arrows indicate examples of PilY1 localized to the tips of sheared T4P filaments. Scale bar, 2 µm.

**Figure S9.**
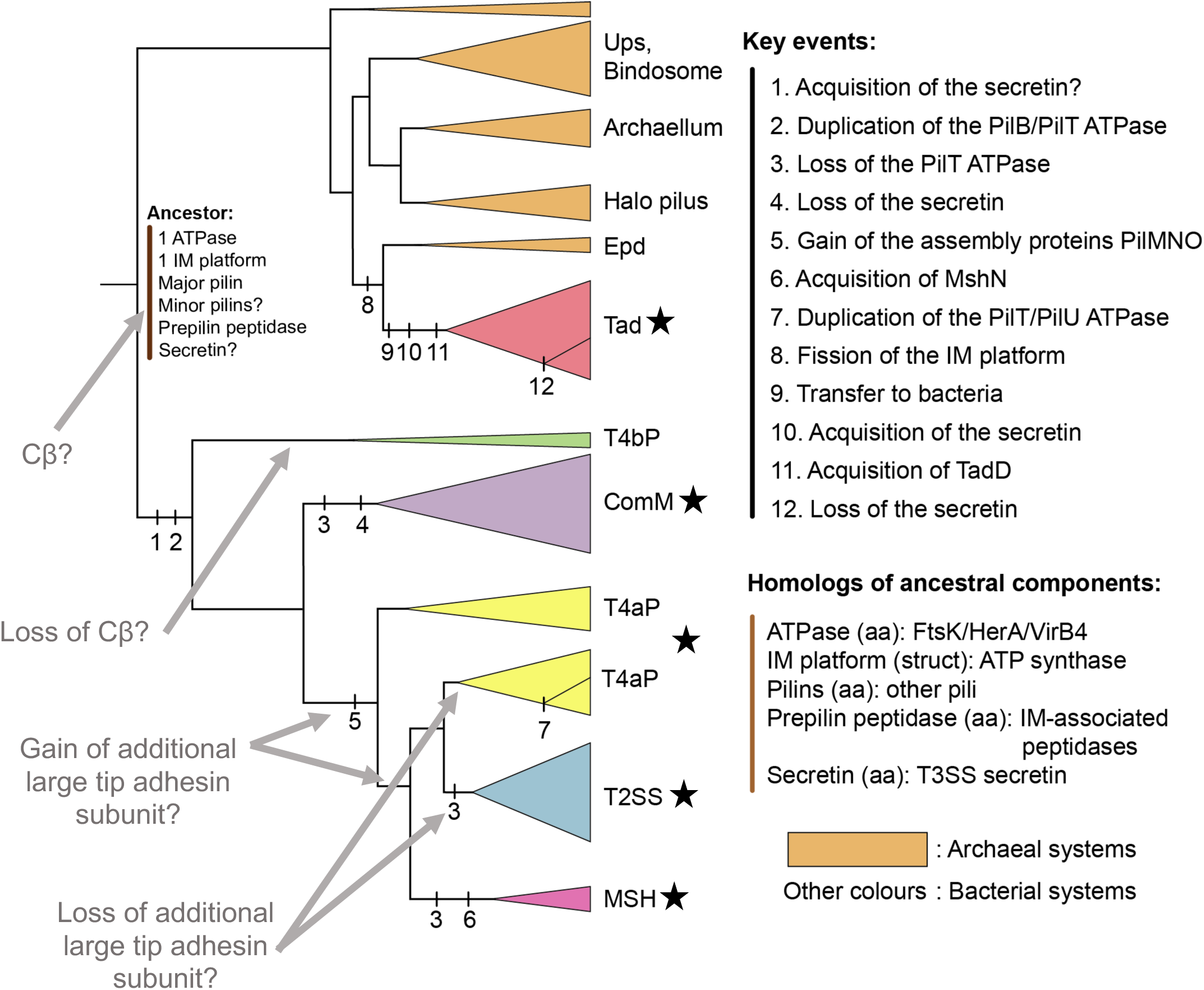
Previously published phylogenetic tree (Figure 7 from Denise et al. 2019) showing the evolutionary relatedness between different types of T4F. Black stars (★) were added here to indicate clades with evidence of β-strand complementation as a mechanism for tip complex licensing. Gray arrows and gray text were added here to indicate potential evolutionary trajectories of tip complex components and aid in discussion. Modified from Denise et al. 2019 Figure 7 legend: The tree was based on the information of the trees of the concatenate and simplified to highlight the key clades and events. The color of the triangles indicates the type of the systems. Each vertical bar on the branch indicates a numbered evolutionary event, whose details are specified under the corresponding number in the list ‘Key events’. The hypotheses for the composition of the last common ancestor of the T4F superfamily are indicated at the root, and the distant homologs of these systems are indicated in the list ‘Homologs of ancestral components’, in which homology was observed by sequence (‘aa’) or structural (‘struct’) similarity. Halo pilus indicates two pili characterised in Halobacteria. Aap, adhesive archaeal pilus; Epd, EppA dependent; IM, integral membrane; MSH, mannose-sensitive hemagglutinin pilus; Tad, tight adherence; T4F, type IV filament; T2SS, type II protein secretion system; T3SS, type III protein secretion system; T4aP, type IVa pilus; T4bP, type IVb pilus; Ups, UV-inducible pilus of *Sulfolobus*.

**Figure S10.**
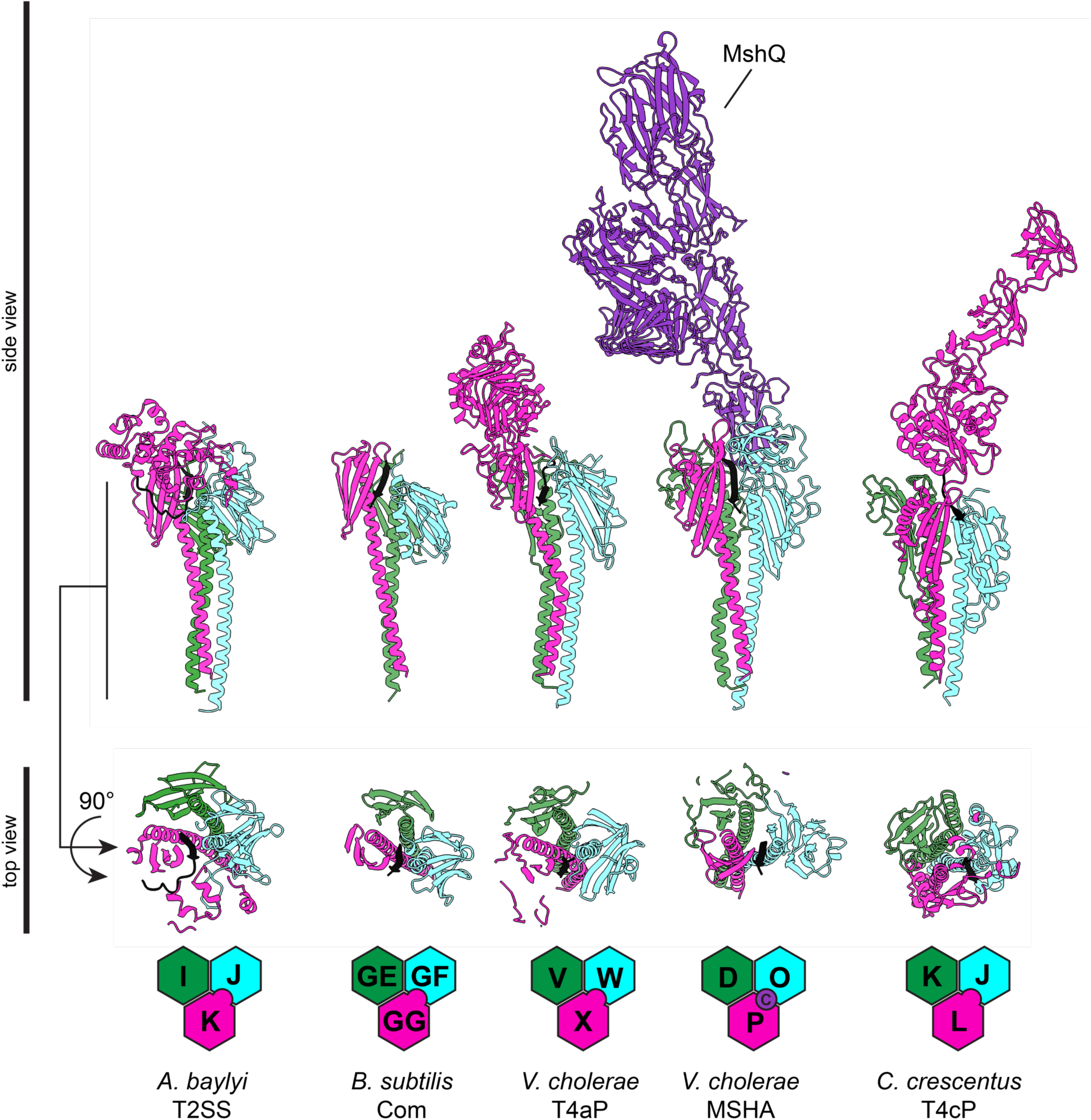
A conserved β-strand is predicted to strand-complement the β-sheet in GspK homologs of diverse T4F. AlphaFold-predicted models of predicted tip complexes for indicated species. In all models, the conserved C-terminal β-strand is colored black.

**Figure S11.**
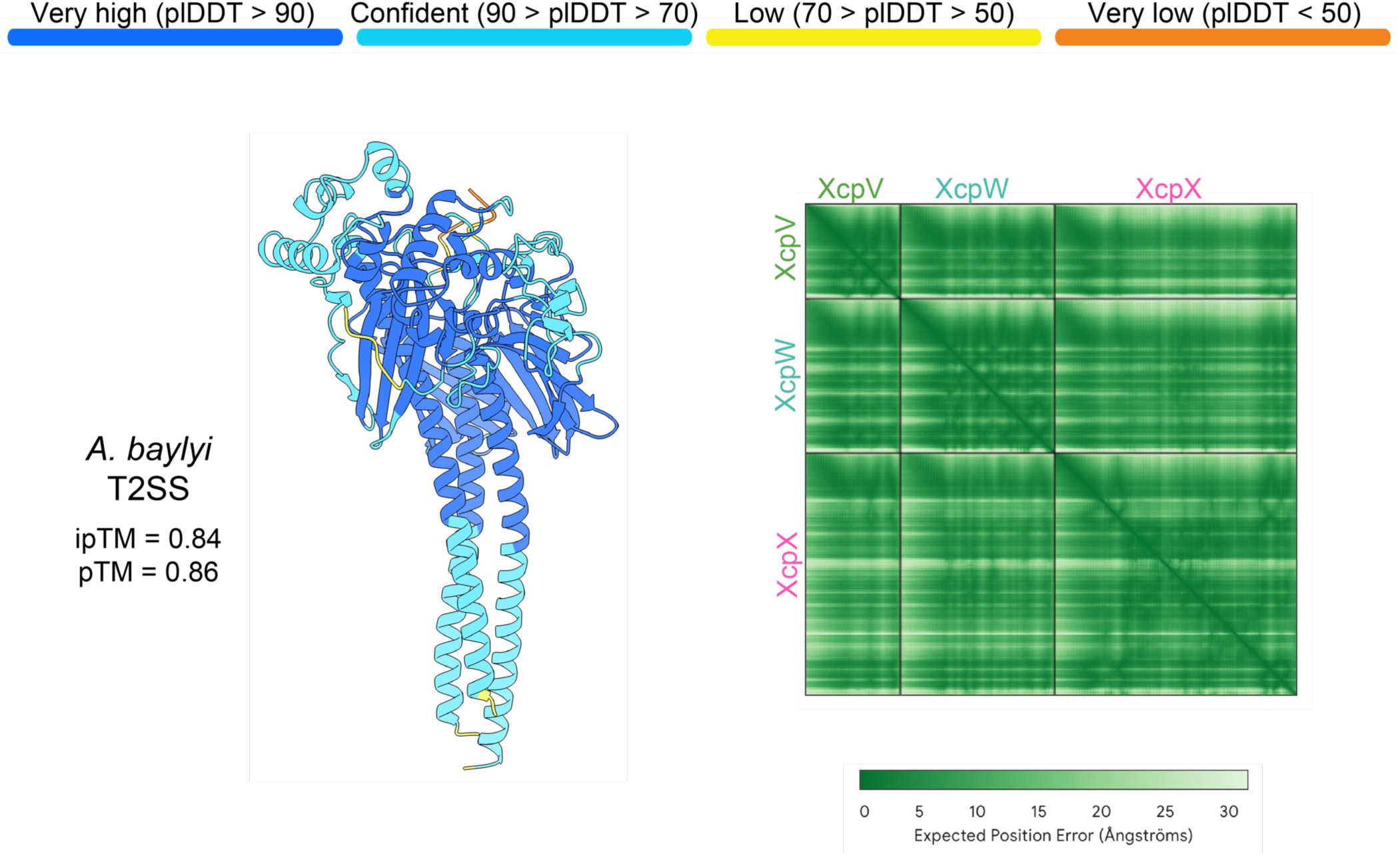
AlphaFold confidence metrics for the tip complex from the T2SS in *A. baylyi*. Structure on the left is colored by confidence intervals indicated at the top. The box on the right shows the predicted aligned error of structure on the left.

**Figure S12.**
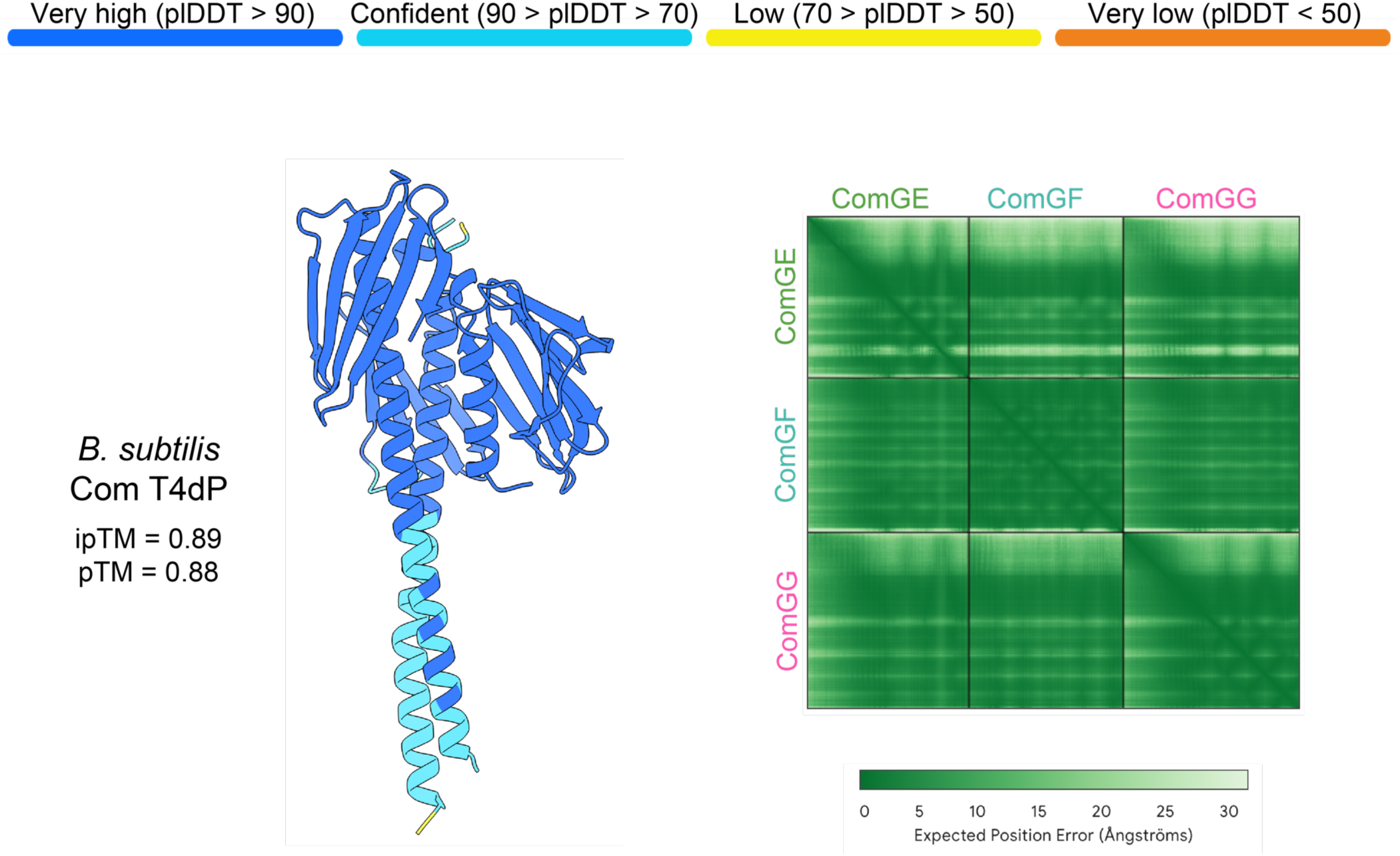
AlphaFold confidence metrics for the tip complex from the Com pilus in *B. subtilis*. Structure on the left is colored by confidence intervals indicated at the top. The box on the right shows the predicted aligned error of structure on the left.

**Figure S13.**
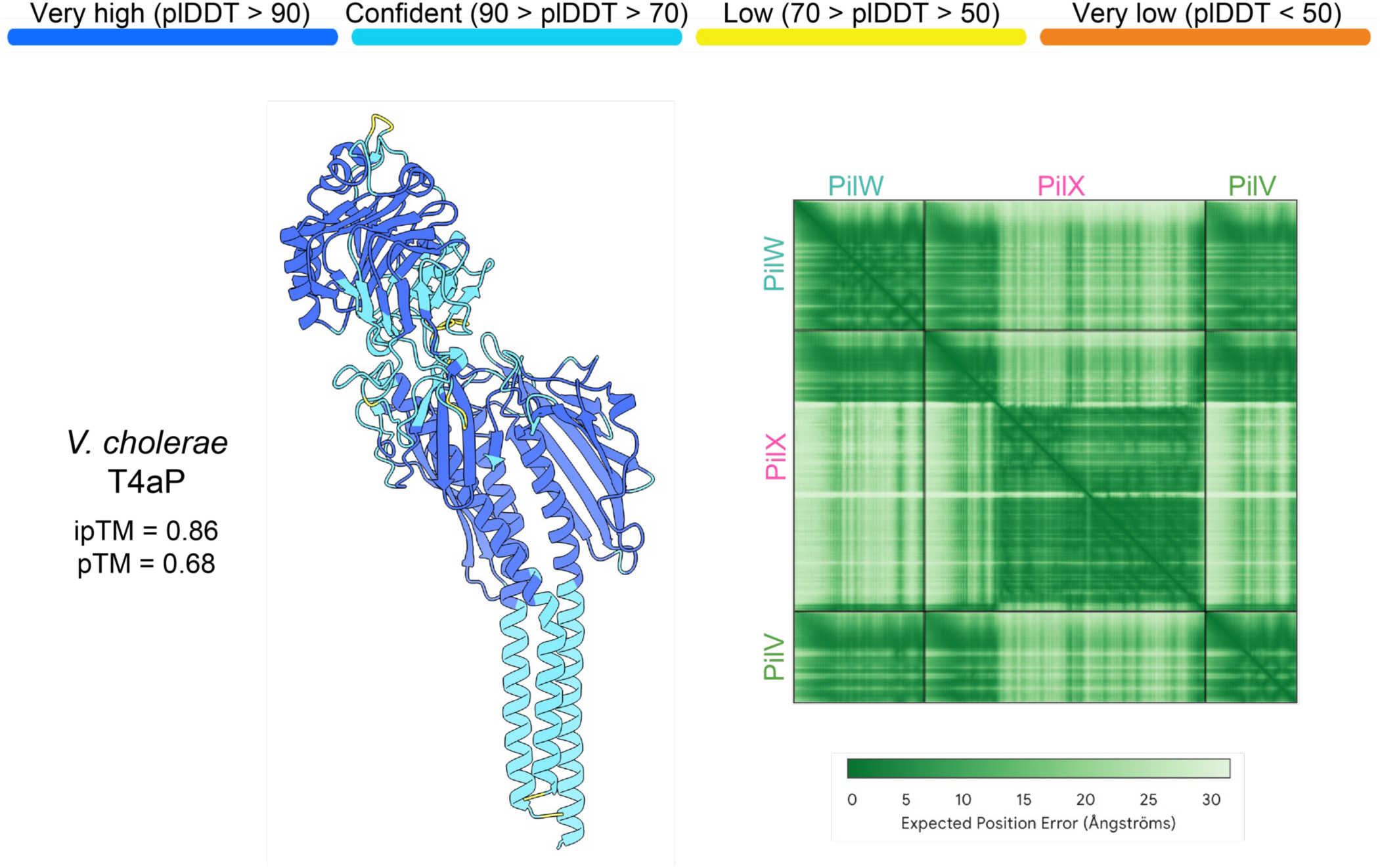
AlphaFold confidence metrics for the tip complex from the T4aP in *V. cholerae*. Structure on the left is colored by confidence intervals indicated at the top. The box on the right shows the predicted aligned error of structure on the left.

**Figure S14.**
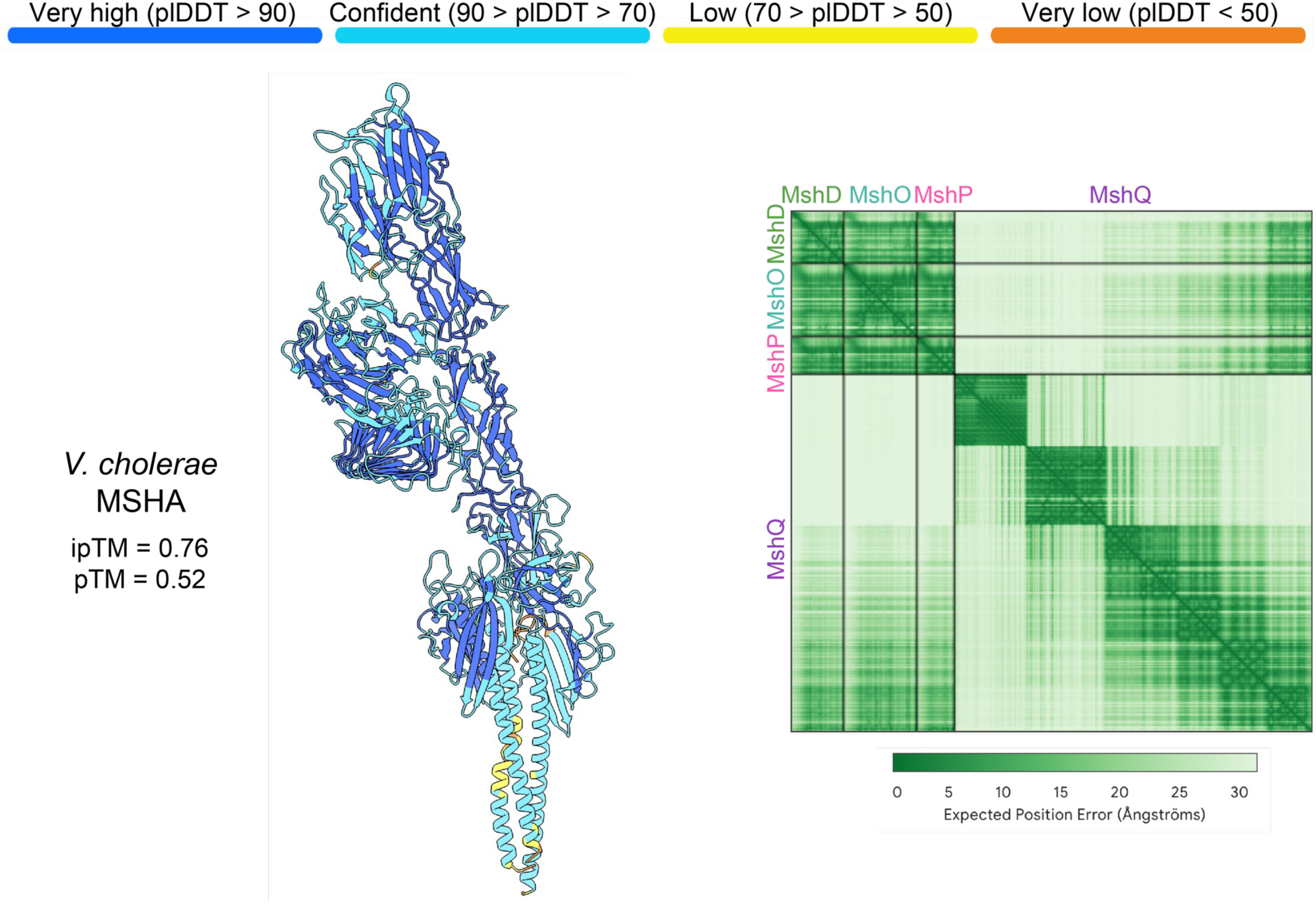
AlphaFold confidence metrics for the tip complex from the MSHA pilus in *V. cholerae*. Structure on the left is colored by confidence intervals indicated at the top. The box on the right shows the predicted aligned error of structure on the left.

**Figure S15.**
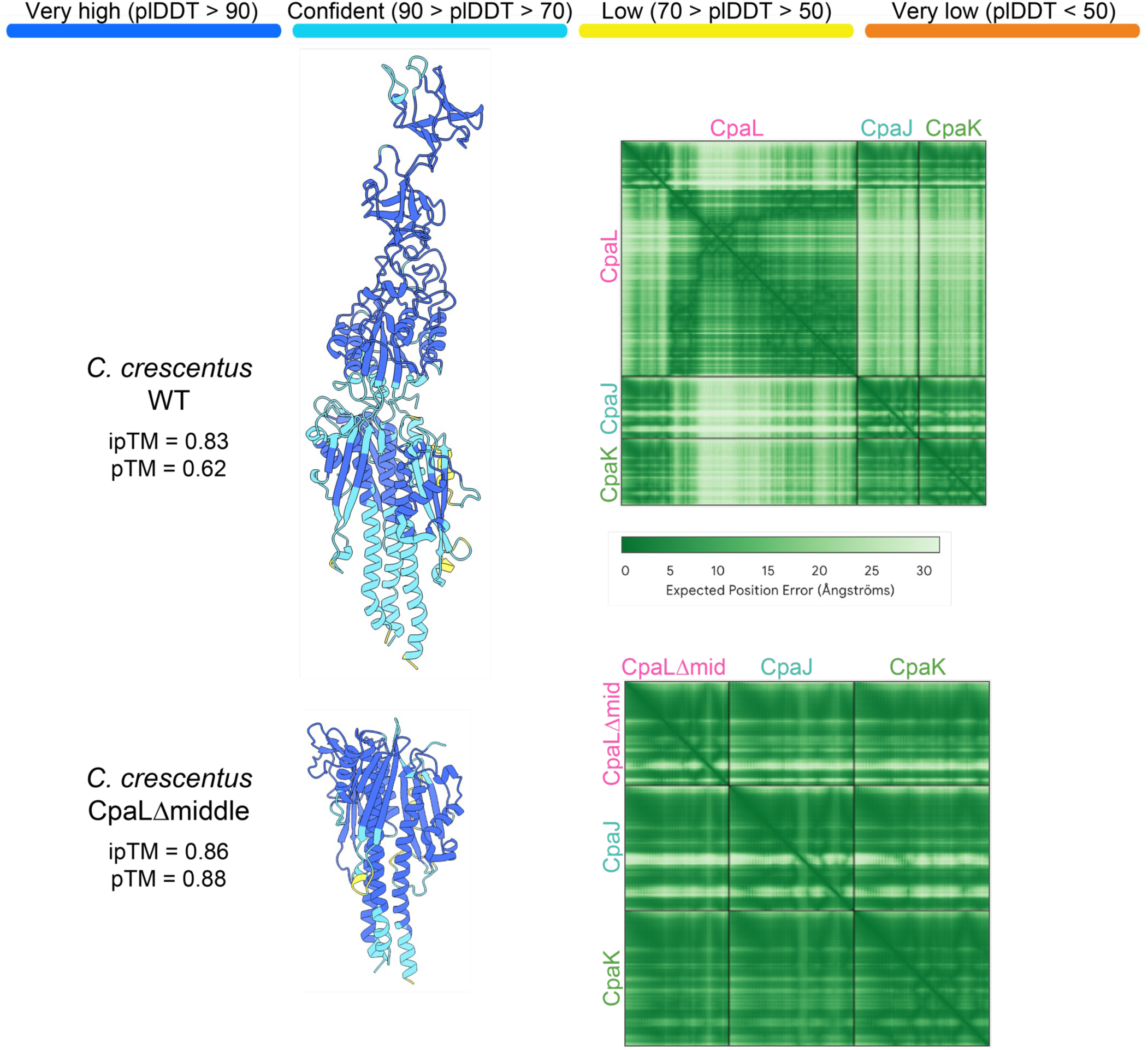
AlphaFold confidence metrics for tip complexes from wildtype T4cP in *C. crescentus* bNY30a and the *cpaL*Δmiddle mutant used in this work. Structures on the left are colored by confidence intervals indicated at the top. The boxes on the right show predicted aligned error of structures on the left.

